# YfiB: An Outer Membrane Protein Involved in the Virulence of *Shigella flexneri*

**DOI:** 10.1101/2021.09.20.461158

**Authors:** Tanuka Sen, Naresh K Verma

## Abstract

The intracellular pathogen *Shigella flexneri*, which is the causative agent of bacillary dysentery, significantly influences the worldwide implication of diarrheal infections, consequentially causing about 1.1 million deaths each year. Due to a non-availability of an authorized vaccine and the upsurge of multidrug resistance amongst *Shigella* strains, there has been a huge demand for further genetic analyses which could help in the advancement of new/improved drugs, and finding vaccine candidates against the pathogen. Whilst many features about the invasion of colonic cells by *Shigella* have been identified, fundamental gaps in information concerning in what way the bacteria transit, survive, and control gene expression, remain. The present study aims to illustrate the role of *yfiB* gene in *Shigella* virulence, which is a part of the periplasmic YfiBNR tripartite signaling system. This system is involved in the regulation of cyclic-di-GMP levels inside the bacterial cells, which is a vital messenger molecule impacting varied cellular processes such as biofilm formation, cytotoxicity, motility, synthesis of exopolysaccharide, and other virulence mechanisms like adhesion and invasion of the bacteria. Through a combination of genetic, biochemical, and virulence assays, we show how knocking out the *yfiB* gene can disrupt the entire YfiBNR system and affect biofilm formation, bacterial invasion, host-surface attachment, and the overall virulence of *Shigella*. This study eventually improves our understanding of the *in-vivo* persistence and survival of *Shigella*, brings light to the c-di-GMP lead regulation of *Shigella* virulence, and provides a prospective new target to design anti-infective drugs and vaccines against *S. flexneri* and other bacterial pathogens.

## INTRODUCTION

*Shigella* is responsible for instigating an invasive enteric infection in humans known as shigellosis and it does this by aggressively attacking the colonic epithelium [1]. It’s a gram-negative bacteria and a well-studied intracellular pathogen, belonging to the *Enterobacteriaceae* family, closely resembling *E. coli* [1, 2]. Globally, shigellosis leads to an enormous amount of morbidity and mortality annually, especially in developing and under-developed countries; significantly contributing to the worldwide burden of diarrheal infections [3,4,5]. Out of the four known *Shigella* species, *S. flexneri* accounts for about 10% of all the diarrhoetic incidents amongst children below 5 years of age in areas of low socioeconomic conditions [6]. In the absence of a globally effective licensed vaccine against *Shigella* and the current treatment method majorly depending on antimicrobial therapy, which keeps getting compromised due to many factors like poor sanitation and hygiene, treatment costs, overuse and/or misuse of antibiotics; there is an ever-growing challenge of antimicrobial resistance in *Shigella*. [7,8,9].

The intracellular lifestyle of *Shigella* including its invasion process and the immune response it generates in the host has been comprehensively studied using various *in-vivo* and *in-vitro* assays, which has expanded our understanding of the pathogen [10]. The virulence genes present on the 30 kb entry region of the large virulence plasmid and the chromosomal genes part of the various pathogenicity islands (PAI) play a crucial role in the survival and infection process of *Shigella flexneri* [1,10,11]. *Shigella*’s pathogenic capability is associated with its ability to adhere, invade, multiply, and eventually kill the colonic epithelial cells [11, 12]. Virulence plasmid-encoded genes such as the type III secretion system (T3SS), *ipA* genes, the *mxi-spa* locus, and chromosomally encode *icsA/virG* gene, have shown to be involved in the Shigella’s survival, adherence, and the invasion process [12–18]. *S. flexneri* while in the host gastrointestinal tract encounters various stimuli and pressures such as the acidic pH, presence of bactericidal bile, competition against host microflora, the compulsion of an effective adherence to the epithelial cells to prevent clearance, followed by a successful invasion and evasion from the host immune system [10,11,19]. To effectively adapt to these adverse conditions in the host during the infection process, intestinal pathogens such as *S . flexneri* have developed and adapted various methods to sense and appropriately respond to external stimuli [20]. One such abundant, highly conserved molecule that acts as a secondary messenger in *Shigella* and other bacteria facilitating these cellular and molecular adaptations to the host environment is Bis-(3’-5’)-cyclic dimeric GMP (c-di-GMP) [20].

The rise in c-di-GMP levels inside the bacterial cell is known to promote flagella/pili biosynthesis, motility, biofilm formation, transcription, exopolysaccharide (EPS) production, surface adherence, invasion of host cells, and overall virulence of the bacteria [21–25]. Inversely a fall in c-di-GMP level can negatively affect all these above-mentioned virulence factors. This association between increased c-di-GMP levels and biofilm production and/or motility mechanisms has been extensively researched in *Escherichia coli*, *Salmonella enterica* serovar Typhimurium, and *Pseudomonas aeruginosa* [22]. Escalated levels of c-di-GMP control biofilm-linked targets such as exopolysaccharide production, bacterial motility, expression of surface adhesin, production of secondary metabolites which eventually help the bacteria in establishing biofilms, effective stress responses, and aids in resistance to antimicrobials [21–27]. The increase in motility due to c-di-GMP regulation, achieved by inducing the formation of pilli/flagella/fimbriae, helps in the formation of biofilms, its dispersal, and maturation leading to a stable three-dimensional biofilm architecture [26, 27]. Additionally, c-di-GMP is involved in adherence/attachment to host surfaces which prevent mechanical clearance and further help in the effective invasion of the host [21–24]. c-di-GMP signaling controls host cell adherence and invasion, intracellular spread, cytotoxicity, regulation of immune responses, and secretion of other virulence factors [21–24]. Therefore understanding how the levels of c-di-GMP are modulated and how this multistep signaling works in *S. flexneri* is essential for developing novel and efficacious treatments against the pathogenic bacteria.

The key protein in question for this study-YfiB, present in the outer membrane and acts as a sensor protein [28]. Complete functional understanding of YfiB remains incomplete, although it is predicted to transduce stress signals into a prompt escalation of intracellular c-di-GMP levels, exopolysaccharide production, and affects downstream virulence factors such as inducing biofilm production [28–30]. YfiB is part of the tripartite signaling system known as YfiBNR, first identified in *Pseudomonas aerouginosa* [31]; which modulates c-di-GMP levels as per the stress indicators detected at the periplasm [28–30]. This signaling system has also been reported in other gram-negatives like *Klebsiella, E. coli*, and *Yersinia pestis* [31]. This system comprises of three different protein members-1) an inner membrane diguanylate cyclase (DGC) responsible for the production of c-di-GMP annotated as YfiN [31, 32]; 2) an outer membrane protein YfiB [31, 32]; and 3) a periplasmic protein YfiN, which forms an association between YfiN and YfiB [31, 32]. From the studies in *Pseudomonas*, it is hypothesized that YfiR allosterically inhibits YfiN by binding to its periplasmic domain and on the other hand YfiB positively regulates DGC activity by sequestering YfiR to the outer membrane, possible via membrane anchoring and peptidoglycan binding; preventing its binding to YfiN and releasing its inhibition [28–32]. This leads to the stimulation of YfiN’s DGC activity, c-di-GMP production, resulting in enhanced biofilm and various other virulence factors [28–32]. The working model for YfiBNR functioning adapted from the *P. aeruginosa* YfiBNR system, developed by Malone, J. G. et al (2010) [31], has been illustrated in Figure 1.

**Figure 1.**
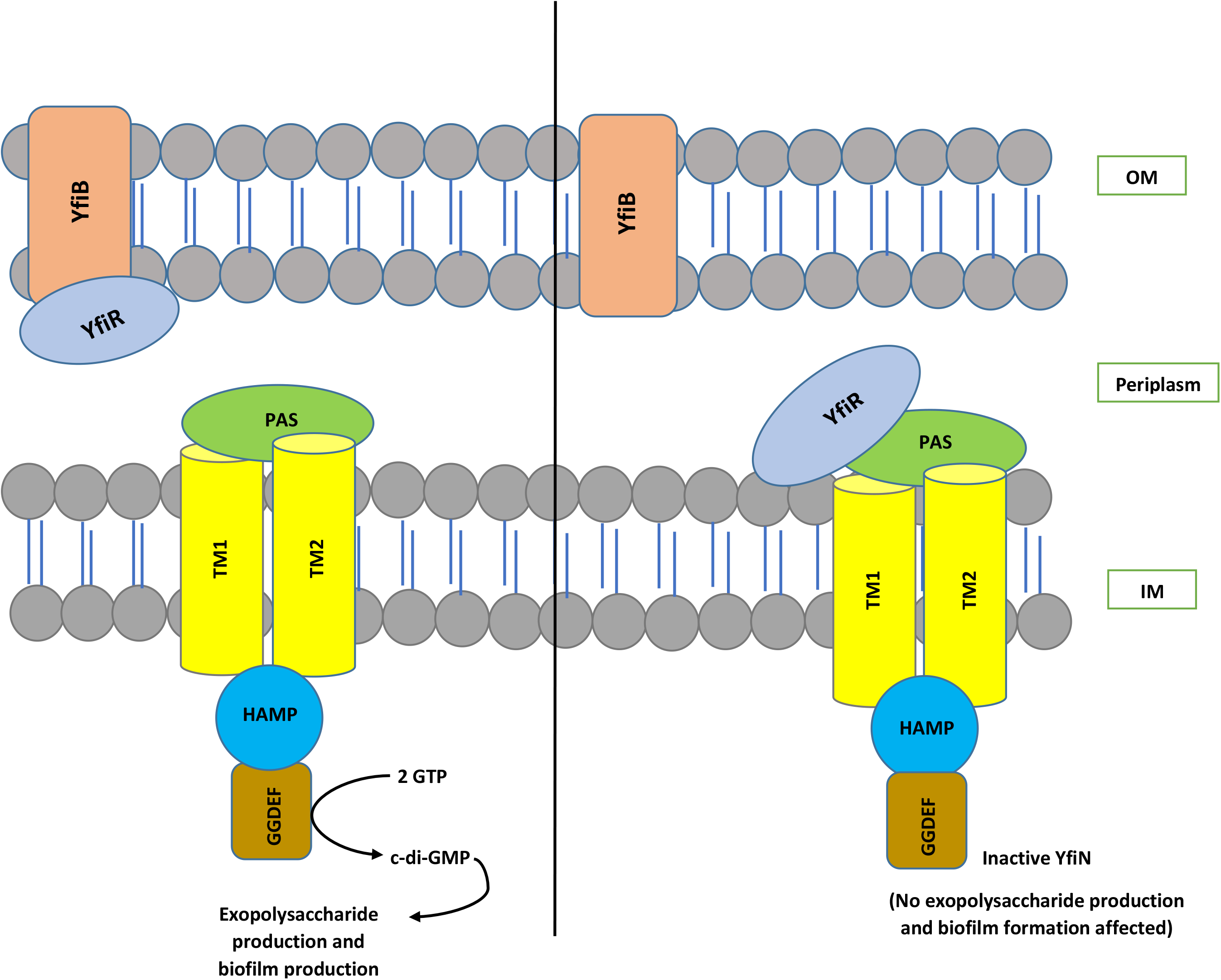
The organization and interaction model of the YfiBNR system. YfiN, an inner-membrane-located DGC is repressed when bound to periplasmic YfiR. While dissociation of the complex by YfiB, which is an Pal-like protein located in the outer-membrane, sequestering YfiR and consequently stimulates YfiN and leads to the production of c-di-GMP.

In this study, we examine the functional role of YfiB in the YfiBNR system and its effect on c-di-GMP production, which consequentially affects many of the virulence factors in *Shigella flexneri*, hence determining the link between YfiN and its positive regulator YfiB. To do this, we generated a yfiB gene knockout, using double homologous recombination and studied its effect on *Shigella*’s survival and pathogenesis, using various *in-vivo* and *in-vitro* virulence assays. We also studied the protein structure in comparison to the YfiB protein of *Pseudomonas*, which previously has been crystallized [30]. This helped in designing targeted mutagenesis experiments to establish which essential amino acids are involved in the functioning of YfiB protein in *Shigella*. This is the first known report to validate YfiB regulation of YfiN’s DGC activity and thereby regulating c-di-GMP levels appropriately is indispensable for *S. flexneri* to cause an effective infection in the host.

## MATERIALS AND METHODS

### Strains and growth conditions

*S. flexneri* serotype 1c (Y394) is a clinical strain from Bangladesh and was generously provided by Nils I. A. Carlin [33]. Using a sterile loop, Y394 and other *Shigella* strains were streaked from glycerol stocks and cultured on Luria Bertani (LB) agar plates and/or tryptic soy broth (TSB) agar plates with 0.01% (wt/vol) Congo red (CR) and were grown aerobically (180 rpm) at 30°C in LB broth to retain the virulence plasmid. Overnight cultures were then sub-cultured at 1:100 dilution and grown at 37°C with 180 rpm shaking, till an optical density of 0.5 to 0.8 (log-phage culture) was obtained at 600 nm (OD600). The antibiotics were added as supplements where indicated at the concentrations: 50 µg/mL erythromycin, 25 µg/mL chloramphenicol, 100 µg/mL ampicillin, 50 µg/mL kanamycin, and 20 µg/mL gentamicin.0.2% arabinose was added for all inducible plasmids. All plasmids, strains, and primers used in this study are listed in **Supplementary Tables S1 and S2.**

### Knocking out yfiB gene to create the deletion mutant and its complementation

*yfiB* deletion mutant (SFL2641/ΔYfiB) was created employing the lambda red double recombination approach established by Datsenko and Wanner with a few modifications [34, 35] using the helper plasmid pKD46, and pKD3 plasmid containing chloramphenicol resistance (CM) for construction of the knockout template. A PCR-based methodology was used for the construction of the knockout template. The CM gene was amplified using the miniprep DNA of pKD3 using primer pairs containing 80bp overhangs, which were homologous to the downstream and upstream of the *yfiB* gene. *S. flexneri* 1c strain (Y394) containing the pKD46 plasmid, encoding the lambda red genes (*gam, beta, and exo*) when induced with 100mM arabinose, was transformed with the knockout template. The positive mutants were first screened on LBA plates containing chloramphenicol and then via colony PCR followed by sequencing.

Complementation of *yfiB* deletion mutant was obtained by cloning the PCR-amplified yfiB gene from the genome of *S. flexneri* 1c SFL1613/Y394 strain, this PCR product was then purified and digested with *Nco*I and *Hind*III. Digested PCR product encoding *yfiB*, was cloned into the pBAD_Myc_HisA vector, which contains a C-terminal 6xHis tag fused to the protein for checking expression (SFL2642/YfiBComp).

### Biofilm Assay

The protocol used for the biofilm assay was tailored from the method initially established by Christensen et al. with a few modifications [19, 36]. A single colony of each bacterial strain was inoculated into a tube containing 10 ml of tryptone Soy broth (TSB). The cultures were incubated overnight at 30°C with 180 rpm shaking. On the following day, each of the cultures was diluted in 1:100 in 5 ml into two tubes one with glucose (1%) and Bile salts (0.4%) and the other without the glucose and bile salts [19, 37, 38]. The OD600 of diluted cultures were checked to make sure the cultures were uniform. 100 µl control media and diluted cultures were inoculated per well in the sterile 96-well microplates (Thermo Fisher). The plates were incubated for 6, 12, and 24 hours at 37°C statically. After incubation, the culture media were removed from the plate gently by slanting the plate and gradually removing the medium using a multichannel pipette, and 1x PBS was used to rinse the wells. The wells were let to dry for 30 minutes, after which methanol was used to fix the dried surface biofilm and then 0.5% crystal violet was used for staining. After staining, the wells were washed 4 times with MilliQ water, followed by drying the plate for over 3 hours. To each well, 200 µl of 95% ethanol was added after drying and incubated for an hour at 4°C to avoid the evaporation of ethanol. The absorbance of all the wells was recorded at 595 nm wavelength (OD595) using the Tecan plate reader. Biofilm Assay with each strain was performed at least 4 times independently with three biological repeats, shown in the graph is the mean of all experiments.

### *in-vitro* bacterial adhesion assay

A bacterial adhesion assay was carried out using Baby Hamster Kidney fibroblasts (BHK) cells [39, 40, 41]. *Shigella* strains were grown in TSB-Sodium deoxycholate medium until an OD600 = 0.6-0.7 (late log phase) was obtained and appropriately diluted in normal saline to a required 2×10^9^ CFU/ml concentration for infection. This prepared culture was used to infect the BHK cells (100 ul/each well), then incubated at 37°C for 2-3 hours for the infection process. After incubation, the cells were rinsed 4 times using 1x PBS, to eliminate all non-adhered bacteria and were lysed using 1% Triton-X by incubating 10 minutes at room temperature on a shaker. LB medium was added to the lysed cells to recover the adhered and invading bacteria. Serial dilutions were prepared of the suspension, plated on LBA, and plates incubated at 37°C, overnight. After incubation, counting of colonies was done and to calculate the total colony-forming unit (CFU). The percentage of adhered bacteria was estimated by dividing the total CFU of adhered bacteria by the total CFU of the inoculum [39].

### *in-vitro* invasion assay and microscopy

The bacterial invasion assays [40, 42, 43] were performed using an epithelial cell line, Baby Hamster Kidney fibroblasts (BHK) cells. Cells were grown in tissue culture flasks (25 cm^2^) to 80% confluency. The incubation of the cells was performed using a CO2 incubator with 5% CO2 at 37°C using the tissue culture media Dulbecco’s Modified Eagle’s Medium (DMEM) supplemented with 10% (v/v) Fetal Bovine Serum (FBS), 2mM Glutamine, and 1x non-essential Amino Acids. Preparing the bacterial culture for invasion assay was done by taking a single isolated colony from each of the *S. flexneri* strains that need to be tested, these were cultured in 5 ml of LB having suitable antibiotics. Cultures were grown overnight at 30°C to preserve the virulence plasmid. Following day, the cultures were diluted 1:100 in LB with 0.1% deoxycholate, containing appropriate antibiotics, and allowed to grow until OD600=0.6-0.8 (mid-log phase) was obtained at 37°C. The appropriate amount of bacteria was harvested via centrifugation and resuspended in 1 ml of 1 x PBS to obtain the culture of 2 x 10^9^ CFU/ml. From this culture, 0.1 ml was used per well of a 6-well tissue culture plate to infect the BHK cells. Centrifugation of the plates for 10 min at 1000X g was done and then incubated for 30 minutes at 37 °C in 5% CO2 level. The media was then removed and wells were washed 4X with 1 ml of 1X PBS. The tissue culture media containing gentamycin (10 µg/ml) was added and incubation of the plate was done for further 60 – 90 minutes (2 hours maximum including 30 minutes above) at 37°C and 5% CO2.

#### Staining and Data collection

After incubation, the media was aspirated and wells were rinsed with 1 ml of 1 x PBS two times. 0.5 ml of filtered Wright-Giemsa stain was added to each well for staining the cells. After 2 minutes, the stain was washed two times with 2 ml MilliQ water and the plates inverted for drying. The invasion of bacteria per BHK cells was determined by enumeration of at least 300 BHK cells [40] and the total bacteria present using a 100X oil immersion microscope, it was performed in duplicates with 3 repeats.

### *in-vitro* Plaque Assay

Plaque assay was carried out using the HeLa cell line [44–45], *Shigella* strains were grown in TSB medium until OD600 = 0.4-0.5 (mid-log phase) was obtained, and appropriately diluted in normal saline to a required 1×10^7^ CFU/ml concentration. This prepared culture was further diluted to 10^6^ to 10^4^ CFU/ml, to infect the HeLa cells (100 ul/each well), and subsequently, the plates were incubated at 37°C, 45-90 minutes for the infection process. The medium was removed after incubation, cells were washed twice with 1X PBS, and 2X gentamycin medium was added to each well. Gentamycin medium was used to remove all external bacteria, which results from futile spreading into the extracellular medium, rather than the neighboring cells. This ensures that each plaque formed, parallels to a sole invasion occurrence at the start of the experiment [43–45]. The plates were then re-incubated at 37°C for 48-72 hours, plaques were mostly visible after 48 hours. When the plaques are visible, the plate was taken out, washed with 1x PBS, to wash away dead cells as they detach from the plate surface. Wright-Giemsa stain is used to counterstain the intact monolayer to make the plaques evident (clear spaces) against the stained background [44, 45]. The overall number of plaques were counted and the mean number of plaques per bacteria was plotted on the graph. The experiment was carried out for 50 technical repeats with two biological repeats each time. Few of the stained plaques were also observed under the microscope to see how the plaques looked under 40X magnification and if there were any observed morphological changes of the cells near and around the plaques.

### *in-vivo* bacterial Accumulation assay using *Caenorhabditis elegans*

The bacterial accumulation assay was executed using the *C. elegans* N2 strain (WT), these nematodes were cultured on a modified nematode growth medium (mNGM), plates already containing *E. coli* OP50 grown on it [46–48]. L4 stage *C. elegans* of a synchronized population was collected off these E. coli OP50 plates using S-basal buffer and were subjected for 3 hours to 200 µg/ml of gentamycin, to remove any *E. coli* OP50 cells present on the surface of the worms. Then, the worms were meticulously rinsed with S-basal buffer to eliminate any remaining antibiotics. Bacterial strains were grown at 37°C, overnight on mNGM plate to prompt the expression of genes encoded by the virulence plasmid. The washed worms were put onto these plates with the seeded *Shigella* strains and incubated for 24 hours, at 22°C. The next day, 15 worms were picked from each plate, and to anesthetize these worms, they were rinsed meticulously using S-basal containing 1mM sodium azide. The worms were subjected for 3 hours, to 200 µg/ml of gentamycin to remove any bacterial cells present on the surface of the worms. After 3 hours, the worms were rinsed 3X times with S-basal buffer containing 1mM sodium azide. The washed worms were resuspended in S-basal buffer containing 0.1% Triton-X and were lysed mechanically using sterile glass beads. The lysate obtained was serially diluted using 1X PBS, and appropriate dilutions were plated on LB agar plates containing the suitable antibiotics, to obtain the intraluminal bacterial counts, these plates were incubated at 37°C for 16 hours. The number of bacteria per 15 nematodes was determined and the analysis was based on three independent repeats.

### *In-silico* analysis of YfiB protein

YfiB protein is present as a hypothetical protein in the *Shigella* genome, hence the 5 step identification system for a hypothetical protein was applied [49]. Functional analysis by identifying conserved domain was done using NCBI-protein BLAST [50] and conserved domain database [51]; followed by analyzing its physicochemical parameter using Expasy’s ProtParam tool [52]; determining the subcellular localization using CELLO [53], PSORTb [54], and PSLpred [55]; the presence of transmembrane helices using TMHMM [56] and HMMTOP [57] tools, and lastly to determine if they are concerned in virulence of *S. flexneri* using VICMpred [58] and VirulentPred [59].

The protein sequence of *Shigella* Y394’s YfiB was analyzed and compared to *Pseudomonas aeruginosa*’s YfiB. The similarity between the protein sequences and the conserved amino acid residues was evaluated using NCBI-protein BLAST and Clustal-omega sequence alignment tool [60]. I-TASSER (Iterative Threading ASSEmbly Refinement) tool [61, 62] was utilized to predict the three-dimensional structure of YfiB protein of Y394, the already solved crystal structure of *Pseudomonas* YfiB protein [29, 30] was used as a template to work out the 3D structure. Significant amino acids of YfiB protein known from previous studies in *Pseudomonas* [29, 30] were identified, and based on this analysis, targets for mutagenesis in Y394 YfiB protein were selected.

YfiB homologs were identified in various other gram-negative species, protein homologs (E-value<10−4) were aligned using Clustal Omega, and conserved amino acid residues were determined. Sequence conservation at each position was evaluated using the WebLogo tool [63], it creates a stack of symbols, the altitude of the pile at each position signifies the sequence conservation and the height of the symbols denotes the comparative incidence of individual amino acids at that locus [63]. Phylogenetic analysis was also carried out of the YfiB homologs and an illustrative tree based on the multiple sequence alignment of YfiB homolog protein sequences was generated using the MEGA software [64], Maximum Composite Likelihood technique was used to determine the evolutionary distances [64, 65, 66]. UniProt accession numbers of all YfiB protein sequences used for the in-silico analysis have been listed in **Supplementary Table S3.**

### Statistical analysis

Statistical analysis to evaluate the significance in each experiment which was based on at least 3 or more trials was done using GraphPad Prism’s unpaired student t-test (standard cutoff for significance: P < 0.05).

### Site-directed mutagenesis and protein expression

For the targeted mutagenesis experiment, four amino acid targets were selected, these being Cys19Gln20 mutated to Ala19Glu20; Pro22Gln23 mutated to Ala22Glu23; Glu29Gln30 mutated to Ala29Glu30, which are part of the YfiB peptidoglycan linker sequence, and Ser36 mutated to Ala36. Site-directed mutants were synthesized by GenScript USA, cloned into the expression pBAD_Myc_HisA vector, using the *BamH*I and *Apa*I sites. For inspecting the expression of the YfiB protein, *Shigella* strains, containing the pBAD_Myc_HisA_YfiB vector were grown in LB broth medium having 100 µg/mL ampicillin and 50 µg/mL erythromycin at 37°C. Once the OD600 of the cultures reached 0.6, 1mL of the cell culture was pelleted down and suspended in an appropriate volume of protein loading dye containing β-mercaptoethanol as the reducing agent to have an equal concentration of proteins in each sample. This protein suspension was boiled at 100°C for 5 minutes before loading 10 µls of it into the wells of two SDS gels. Coomassie blue staining of one loaded SDS gel was done to check for equal total protein loading and western transfer with blotting was carried out with the second gel using the anti-HisA [67] antibody to check for protein expression (**Supplementary Figure S3A and S3B**).

## RESULTS

### Biofilm formation is significantly slower in the *yfiB* deleted *Shigella* strain

Enteric pathogens like *Shigella* produce biofilms to enhance survival and protect themselves from harsh environments such as the bactericidal bile present in the human GIT [19, 37]. Biofilms are defined as structured groups of bacteria entrenched in a self-constructed matrix that is made up of various proteins, extracellular DNA, and exopolysaccharides [19, 37]. c-di-GMP is known to be involved in regulating many virulence phenotypes in bacteria including biofilm production [26, 27]. To test this hypothesis in *Shigella* and to assess whether the YfiB protein along with the YfiBNR system plays a role in biofilm formation, we knocked out the *yfiB* gene from *S. flexneri* 1c strain (SFL1613/Y394) and measured the extent of biofilm produced by the wild-type and mutant strain (SFL2641/ΔYfiB) at 6, 12, and 24 hours in presence and lack of 0.4% bile salts and 1% glucose [19, 37]. The *yfiB* gene was also complemented in the knockout strain (SFL2642/YfiBComp) by cloning the gene in the pBAD_Myc_HisA vector and introducing this plasmid in the SFL2641 strain, to see if the wildtype function is reinstated.

As seen in **Figure 2A**, biofilm formation remained minimal in the lack of bile salts and glucose, in comparison to a substantial increase of biofilm formation when bile salts and glucose is present. A significantly slower capability to produce biofilm was observed in the *yfiB* deletion mutant (SFL2641/ΔYfiB) when compared to the WT (SFL1613/Y394) strain. The biofilm produced by SFL2641 was considerably affected at 6 hours of incubation but showed no difference to the WT biofilm production at 12 and 24 hours. The complemented strain (SFL2642) showed similar amounts of biofilm production as WT and hence it could be concluded that complementing the *yfiB* gene in the mutant strain can reinstate the wildtype function (**Figure 2B**).

**Figure 2.**
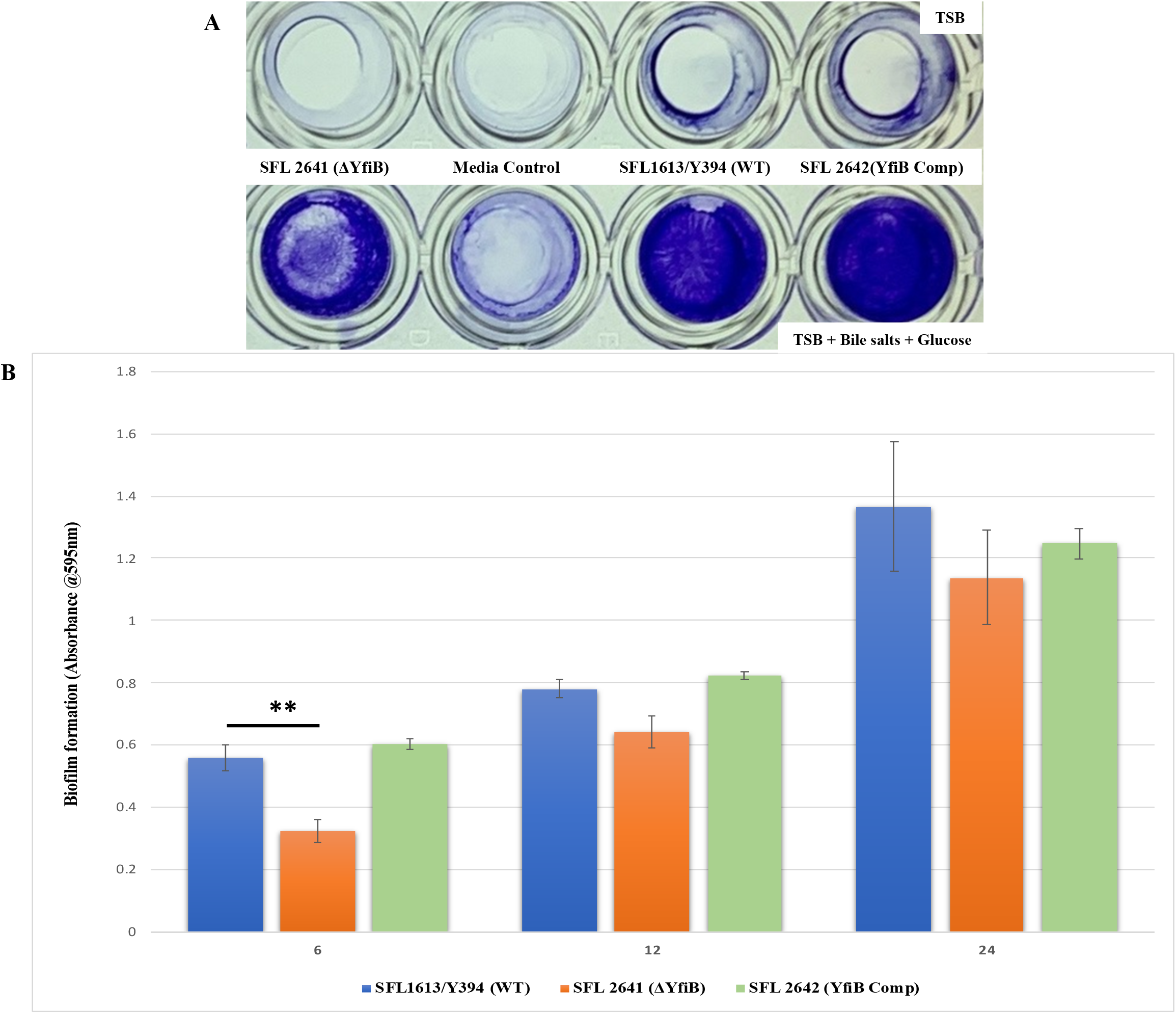
Quantification analysis of biofilm formation by *S. flexneri* strains. **(A)** Examination of biofilm formation in plate wells when grown with and without bile salts in TSB media at 24 hours. Biofilm formation was performed in a 96 well plate, the wild-type *S. flexneri* serotype 1c (SFL1613/Y394), *yfiB* deletion mutant (SFL2641/ΔYfiB) and *yfiB* complemented strain (SFL2642/YfiBComp) were cultured in tryptone soy broth (TSB) medium with and without bile salts and glucose for 6, 12 and 24 hours static. Observed biofilm was stained using crystal violet and assessed by determining the optical density at 595 nm (OD595). **(B)** Biofilm formation of WT (SFL1613/Y394), *yfiB* deletion mutant (SFL2641/ΔYfiB) and *yfiB* complemented strain (SFL2642/YfiBComp) at 6, 12, and 24 hours; at 6 hours, considerable difference was seen between the WT strain and *yfiB* deletion mutant. The Y-axis represents the biofilm amount relative to the blank media. A Student’s t-test was applied for pairwise comparison of the biofilm formation and the analysis was based on three independent biological repeats with each assay having at least four technical repeats. The asterisks represent the variation observed is statistically significant (p<0.05) and error bars depict the SD of data based on the three independent experiments.

### *yfiB* deleted *Shigella* strain shows a decreased adhesion and invasion of BHK cells and displays an attenuated ability to form plaques in HeLa cells

Infiltrating the colonic and rectal epithelium, proliferating intracellularly, spreading from cell to cell, are the three fundamental features of all virulent *S. flexneri* strains [68], which characteristically leads to severe host tissue damage and other shigellosis associated manifestations [1]. Initial attachment by *Shigella* to the host surface is essential for consequent infection and invasion, and how this is achieved, is a process that still needs further research. In recent studies virulence factors such as *icsA/virG*, which is a surface-displayed protein was discovered to play a crucial role in *Shigella*’s host cell adhesion [16]. In another study, it was observed that *Shigella* expressed genes of long polar fimbriae, Type 1 fimbriae, and of the curli operon which helps the bacteria in adhesion to host surfaces and biofilm production [18]. Various attachment and internalization mechanisms are required by *Shigella* for the invasion of mammalian cells, with the bacteria eventually residing in the host cell cytoplasm [11, 12, 69]. After the invasion, the intercellular spread can happen in two ways-1) lysis of the infected cells and discharge of intracellular bacteria; or 2) individual bacteria evading the initially infected cell before lysis and attacking the neighboring cells without being exposed to the outside environment [44]. Even in the second case, the initially infected cell is killed or lysed by the intracellular bacteria [44]. This kind of interaction proposes that an individual *Shigella*-infected host cell would lead to an advanced infection of the surrounding cells and leave behind a region of dying or dead host cells, which is also known as a plaque [44, 45]. To determine the set of virulence genes responsible for the adhesion, invasion, and intercellular spread in host cells, *in-vitro* assays with cultured epithelial cell monolayers such as HeLa and BHK cell lines have been extensively used, which mimics the *in-vivo* infection model [40].

To investigate the involvement of the *yfiB* gene and *yfiBNR* operon in the initial attachment or adherence of bacteria to the host cell, an in-vitro adhesion assay was carried out in which we infected BHK monolayers with the *Shigella* strains-WT (SFL1613/Y394), *yfiB* deletion mutant (SFL2641/ΔYfiB), and *yfiB* complement strain (SFL2642/YfiBComp). The percentage of adhered bacteria of the WT strain (SFL1613/Y394) was seen to be four folds higher than the *yfiB* deletion mutant (SFL2641/ΔYfiB). Statistical analysis via an unpaired student t-test confirmed that the *yfiB* deletion mutant had a considerably decreased capacity to adhere to the epithelial cells of the host when paralleled with the wild-type (**Figure 3**). When the *yfiB* gene was complemented (SFL2642/YfiBComp), it was seen that the wild-type adherence ability was restored. This clearly shows the role of YfiB and YfiBNR system in *Shigella* virulence, as the very first step of its infection process is adherence to host cells.

**Figure 3.**
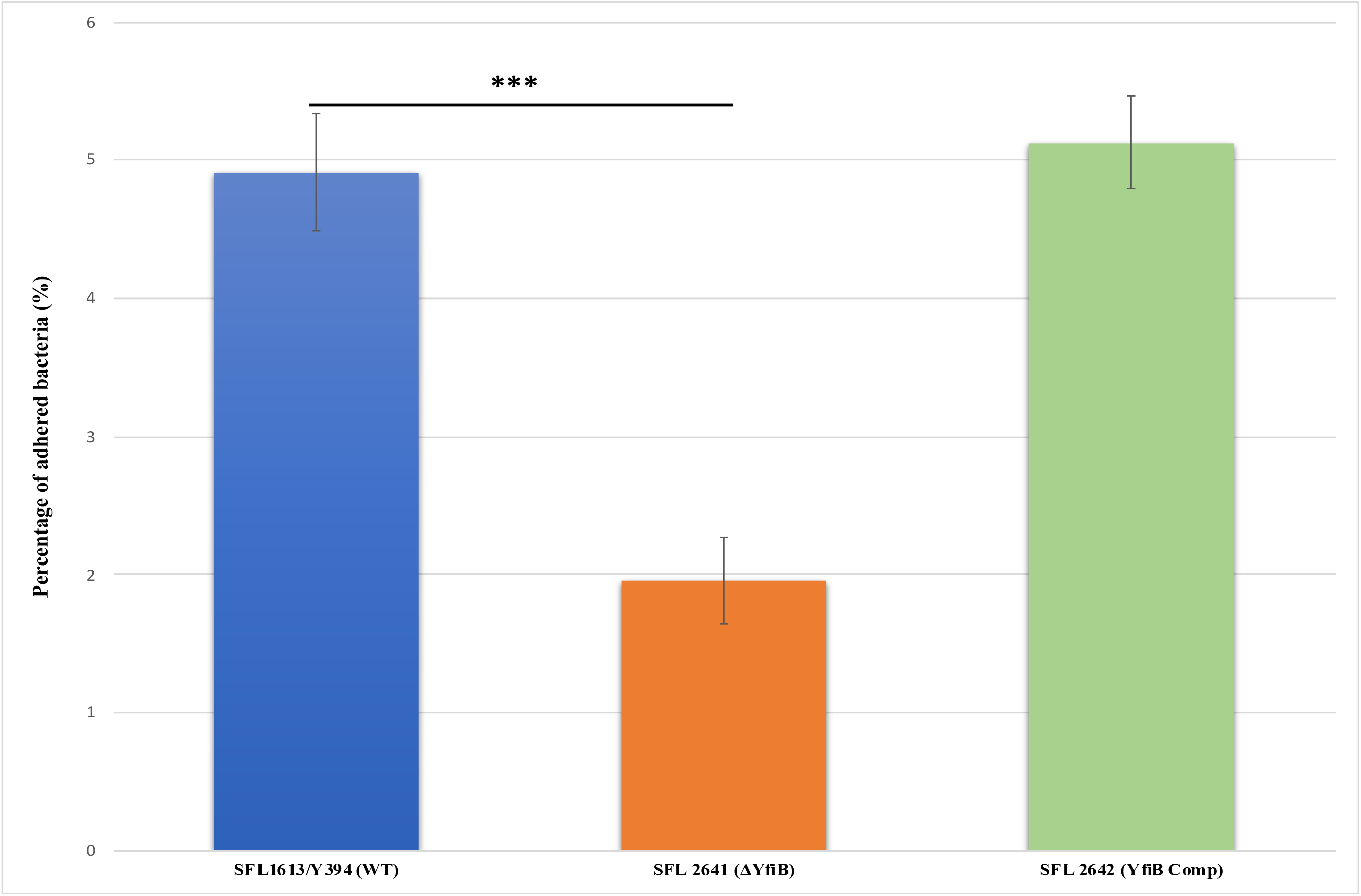
Graphical representation of percentage adhesion to BHK cells by *S. flexneri* strains. The wild-type *S. flexneri* serotype 1c (SFL1613/Y394), *yfiB* deletion mutant (SFL2641/ ΔYfiB) and *yfiB* complemented strain (SFL2642/YfiBComp) were used to infect the BHK cells. After infection, BHK cells were washed and lysed; lysates were serially diluted before plating on an LB agar plate for enumeration and CFU calculation. The percentage of adhered bacteria was calculated by dividing the total colony forming unit (CFU) of adhered bacteria by the total CFU of the inoculum. The Y-axis represents the percentage of adhered bacteria to BHK cells after 2 hours of post-infection at 37°C/ 5% CO2. The results were obtained from six autonomous experiments and to analyse the statistical differences in the adhesion, a student’s t-test was performed. The asterisks represent that the variance observed were statistically significant (p<0.05). SD of data from the six autonomous experiments is depicted by the error bars.

Examining the consequence of deleting the *yfiB* gene and disrupting the *yfiBNR* operon on the invasive potential of *S. flexneri* serotype 1c, was done by carrying out an in-vitro invasion assay, in which BHK cells were infected with the strains-WT (SFL1613/Y394), *yfiB* deletion mutant (SFL2641/ΔYfiB), and *yfiB* complement strain (SFL2642/YfiBComp). The bacteria invading the BHK monolayers were stained and counted by microscopy, the average amount of bacteria invading per BHK cell was found to be 4-folds higher in wildtype when compared with the *yfiB* deletion mutant strain (**Figure 4A**). It was additionally observed that the spread of bacteria to neighboring cells was far less in the deletion mutant when paralleled to the wild type, demonstrating that the deletion affects the intercellular spread of the infection. Statistical analysis confirmed that the *yfiB* deletion mutant had a considerably reduced capability to invade host epithelial cells when paralleled with the wild-type strain. When the *yfiB* gene was complemented, it was seen that the wild-type invasion ability was restored, which indicates the role of YfiB and YfiBNR system in the invasive potential of *S. flexneri* serotype 1c (**Figure 4 B-E**).

**Figure 4.**
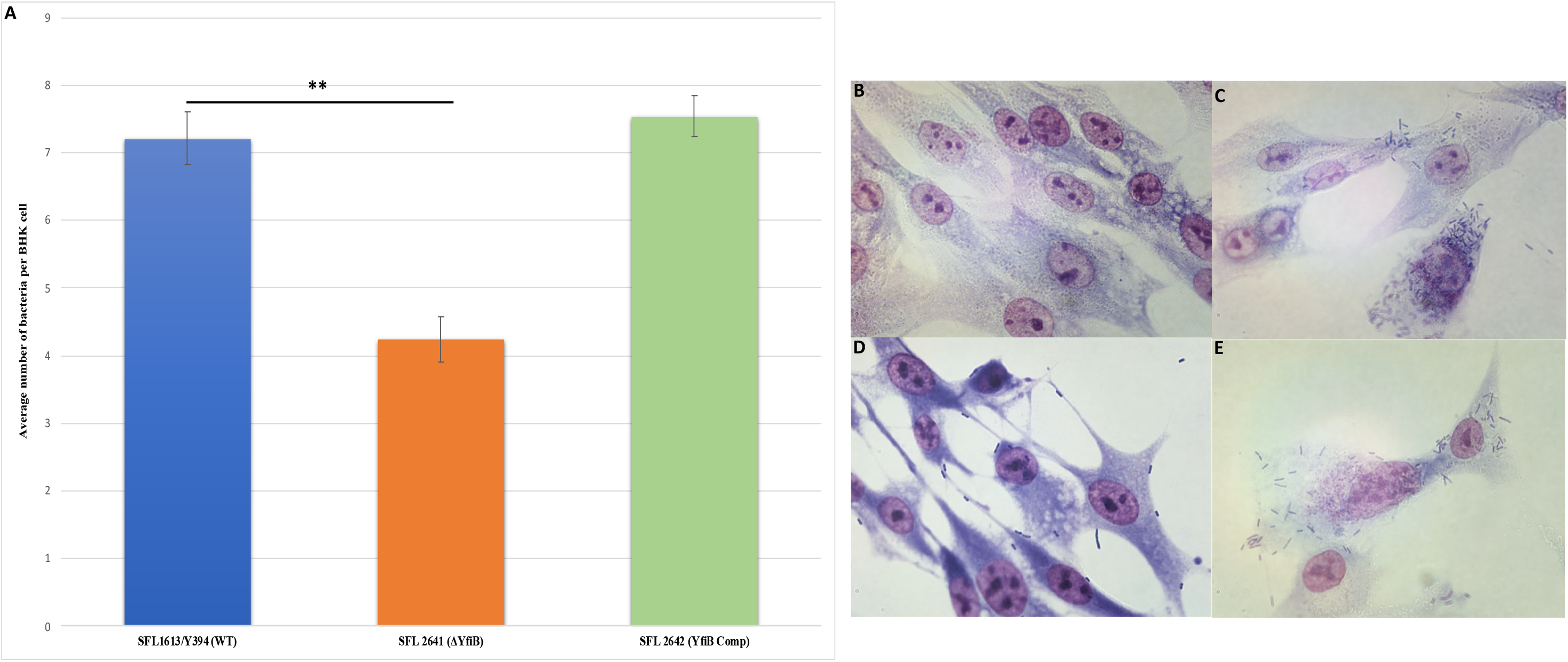
Invasion of BHK cell monolayers by *S. flexneri* strains. **(A)** The wild-type *S. flexneri* serotype 1c (SFL1613/Y394), *yfiB* deletion mutant (SFL2641/ΔYfiB) and *yfiB* complemented strain (SFL2642/YfiBComp) were used to infect the BHK cells. The Y-axis represents the number of internalized bacteria per BHK cell after 2 hours of post-infection at 37°C/ 5% CO2. The results were obtained from three independent experiments and were derived from scoring at least 300 BHK cells. To analyse the statistical differences in the invasion, a student’s t-test was performed. Asterisks represent that the variance observed were statistically significant (p<0.05) and the SD of data from three autonomous experiments is depicted by the error bars. Microscopic image of infected BHK cells, after 2 hours of post-infection at 37°C and at 5% CO2, Giemsa stain was used to stain the monolayers, and the plates were observed under the microscope. **(B)** Microscopic image of non-infected BHK cells under 100X oil immersion. **(C)** Microscopic image of BHK cells infected with the wild-type strain (SFL1613/Y394) under 100X oil immersion**. (D)** Microscopic image of BHK cells infected with *yfiB* deletion mutant (SFL2641/ΔYfiB) under 100X oil immersion. **(E)** Microscopic image of BHK cells infected with *yfiB* complemented strain (SFL2642/YfiBComp) under 100X oil immersion.

A plaque assay was also carried out using HeLa cells to understand the two vital steps involved in the *Shigella* infection process, them being invasion and intracellular spread of the bacteria [44]. The number of plaques evaluates the invasive ability of the bacteria and the size dimensions of the plaques reveals the bacteria’s capability to spread within an epithelium [44, 45]. We infected HeLa cells with *Shigella* strains-WT (SFL1613/Y394), *yfiB* deletion mutant (SFL2641/ΔYfiB), and *yfiB* complement strain (SFL2642/YfiBComp) for 48-72 hours. As seen from **Figure 5A**, which depicts the mean number of plaques produced by each strain, the number of plaques produced by the *yfiB* deletion mutant (SFL2641/ΔYfiB) was 2.5 times lesser than the wild-type strain (SFL1613/Y394). When the *yfiB* gene was complemented (SFL2642/YfiBComp), it was seen that the wild-type plaque-forming ability was restored. There is no substantial difference observed in the diameters of plaques formed between the strains or any other plaque characteristics (**Figure 5B**). Microscopy was also performed to observe the stained HeLa cells for any morphological changes near and around the plaques under 40X (**Figure 5 C & D**). A low number of plaques seen in the *yfiB* deletion mutant indicates its inability to adhere to the host cells and disseminate from the initial site of infection, thereby showing significantly less intercellular spread, compared to the wild type.

**Figure 5.**
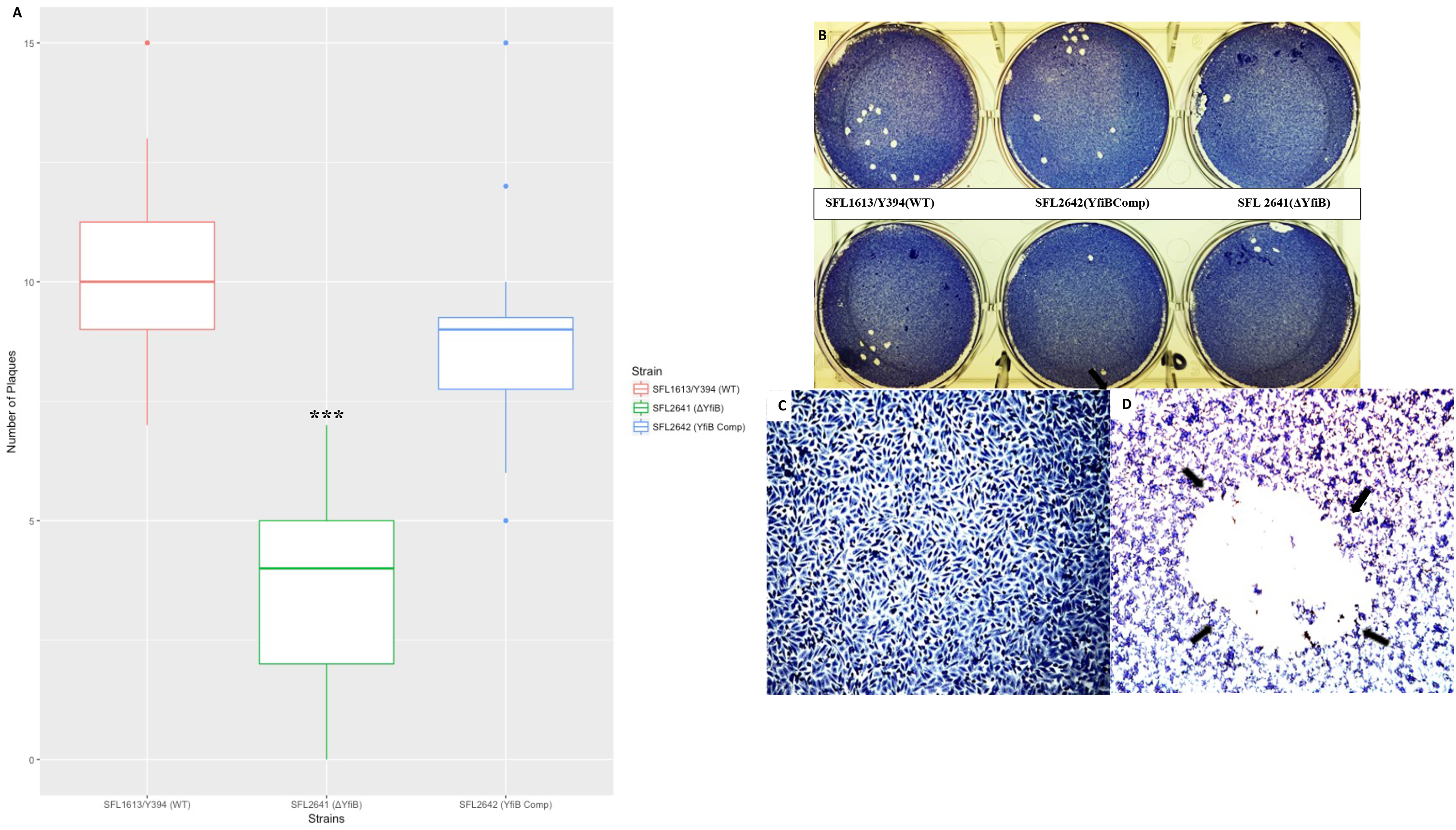
Plaque formation in HeLa cells by *S. flexneri* strains. **(A)** Boxplot showing the mean number of plaques formed by each strain-wild-type (SFL1613/Y394), its isogenic *yfiB* deletion mutant (SFL2641/ΔYfiB) and *yfiB* complemented strain (SFL2642/YfiBComp). Plaque assay was performed in 6-well plates by infecting confluent HeLa cell monolayers with *Shigella* strains. Infection was carried out for 48-72 hours, post-infection, the monolayer was washed and stained with Giemsa, resulting in clear plaques that are visible against the coloured background. The number of plaques formed were counted for each strain and then plotted. The results were obtained from 50 independent technical repeats with two biological repeats each time. To analyse the statistical variances in plaque formation between the strains, a student’s t-test was performed and statistically significant (p<0.05) variations are denoted by the asterisks. **(B)** Visual assessment of the plaques formed on the cellular monolayer at 72 hours post-infection, clear plaques representing dead cells in a well of 6-well plate with HeLa monolayer stained with Giemsa. **(C)** Microscopic view of an uninfected monolayer of HeLa cells stained with Giemsa, under 40X magnification. **(D)** Microscopic view of *Shigella* infected monolayer of HeLa cells stained with Giemsa, under 20X magnification, arrows point to an entire individual plaque.

### *yfiB* deletion influences *C. elegans* lifespan and survival during infection

*Shigella* has a very narrow host range, infecting only primates and humans [1], the most commonly used *in-vivo* animal models are the murine pulmonary and guinea pig keratoconjunctivitis models [46]. Though these animal models have been used for years, they lack clinical relevance of infection site and the symptoms produced when compared to a *S. flexneri* infection in humans [46]. *C. elegans* which is a soil-dwelling nematode recently has been considerably used for several enteric pathogens to study host-pathogen interactions [47]. *C. elegans* show morphological resemblances with the human intestinal cells and significant similarities with the human innate immune system, making it a good animal model to study enteric bacterial infections [46–48]. It was first revealed from the studies carried out by Burton et. al., that there is an accumulation of virulent strains of *S. flexneri* in the *C. elegans* guts, affecting the lifespan of the worms, however, the avirulent strains tend to get digested [48]. Bacterial accumulation of pathogenic bacteria in the worm’s intestine can drastically shorten the lifespan of worms and on the other hand, accumulation of non-pathogenic *E. coli* such as OP50, which is ingested as feed increases the lifespan of worms [46–48]. Thus, bacterial accumulation assay using *C. elegans* assisted us in evaluating the role of *yfiB* gene, the *yfiBNR* operon system, and modulation of c-di-GMP in *Shigella*’s infection of the nematodes and evading its immune defense system.

A synchronized population of L4 larva stage hermaphrodite N2 nematodes was prepared and fed with wild-type (SFL1613/Y394), *yfiB* deletion mutant (SFL 2641/ΔYfiB), and the *yfiB* complemented strain (SFL2642/YfiBComp). Accumulation of the wild-type strain was around two-fold higher (average CFU/worm) than that of the mutant (average CFU/worm for SFL 2641/ΔYfiB). The average CFU/worm for the complemented strain (SFL2642/YfiBComp) was similar to that of the wildtype, indicating that the invading ability of the wild-type strain was restored. Statistical analysis confirmed that the accumulation in the *C. elegans* intestinal lumen by the wild-type strain was considerably higher compared to the *yfiB* deletion mutant (**Figure 6**).

**Figure 6.**
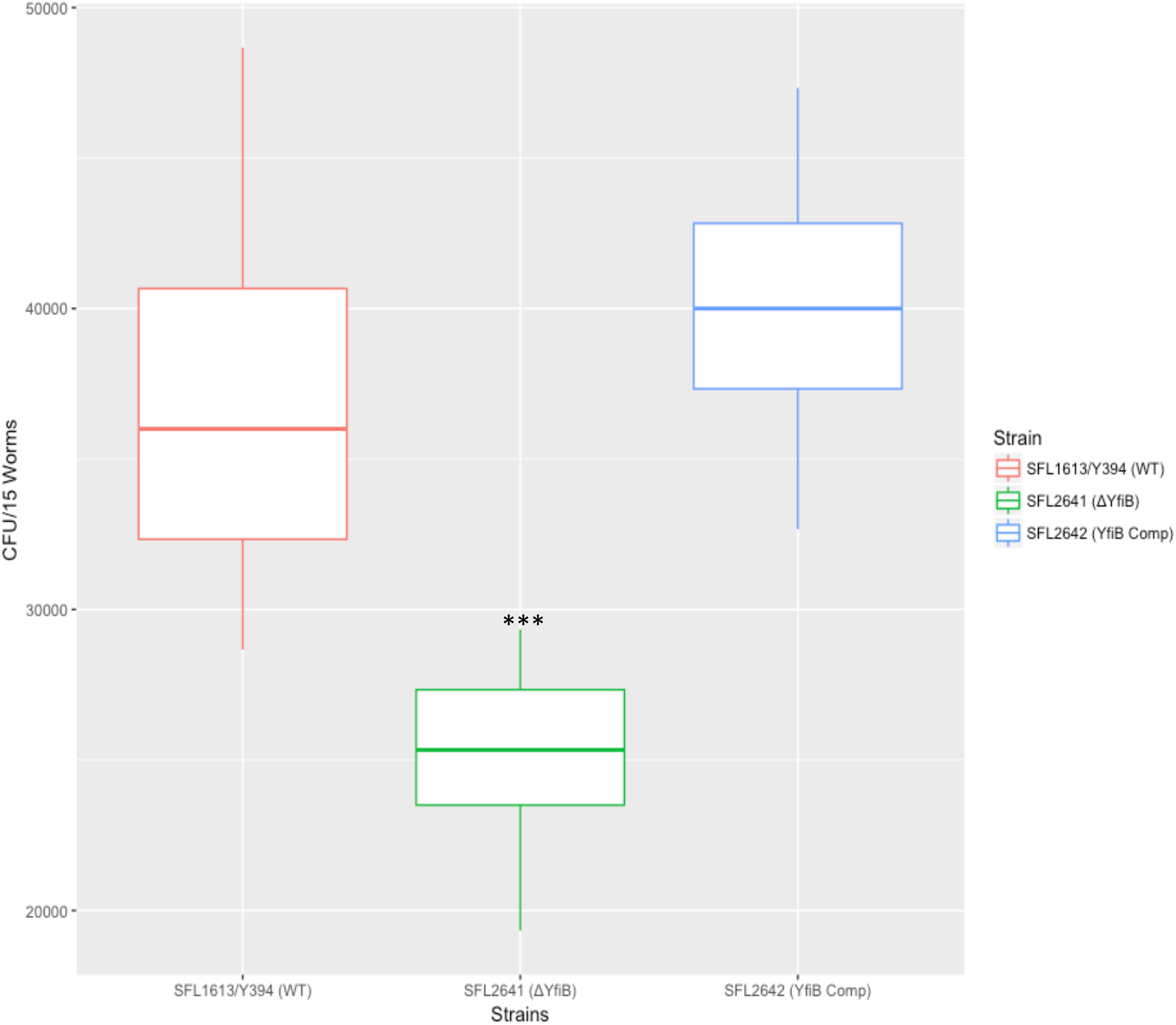
Quantification of bacterial accumulation in *C. elegans.* The synchronized L4 stage N2 nematodes were fed with wild-type (SFL1613/Y394), *yfiB* deletion mutant (SFL2641/ΔYfiB) and *yfiB* complemented strain (SFL2642/YfiBComp) for 24 hours at 22°C. From each plate, 15 worms were picked; lysed utilising glass beads, after which suitable dilutions of individual lysate were plated on LB agar plates to attain bacterial accumulation numbers. To analyse the variance in accumulation seen between the strains, a students t-test was performed. Wild-type strain (SFL1613/Y394) and *yfiB* complemented strain (SFL2642/YfiB Comp) showed higher accumulation in *C. elegans*, when compared to *yfiB* deleted mutant (SFL2641/ΔYfiB) strain. Statistically significant (p. value < 0.05) variance is indicated by the asterisks and the analysis was based on three independent repeats.

### In-silico investigation of the YfiB protein elucidates its structure, function, and conserved nature across bacterial species

YfiB was present as a hypothetical protein in the *S. flexneri* 1c strain (Y394) genome, *in-silico* functional analysis was carried out using NCBI-protein BLAST and the conserved domain database to find out homologs across other bacteria and conserved domains. Homologs across other gram-negative bacterial species were found. YfiB is a 17.2 KDa putative lipoprotein or an integral membrane protein, with an OmpA domain and Omp_C-like superfamily domain (**Supplementary Figure S1A**). Functional prediction of YfiB as per the conserved domains showed that it might be involved in transmembrane transporter activity; cell growth and adherence; in response to stress and/or external encapsulating structure. Physiochemical properties, subcellular location, presence of transmembrane domains, and involvement in virulence were also studied, using various bioinformatics tools listed in the methods section. YfiB was predicted to be located in the outer membrane, comprising of a PAL-like peptidoglycan (PG) domain and no transmembrane helices. Consensus prediction of gene ontology (GO) showed that YfiB was involved in molecular and biological processes of the bacteria and the protein had a 0.271 grand average of hydropathicity (GRAVY).

YfiBNR system has been previously studied in *Pseudomonas* [31, 32], and protein crystal structures of all the three proteins involved in the operon have been solved [28–32]. When the YfiB protein sequence of *Shigella* was compared to that of *Pseudomonas* via NCBI nucleotide and protein BLAST, no considerable similarity was found between the nucleotide or protein sequences as per the default BLAST settings. The protein sequence of YfiB from *Shigella* and *Pseudomonas* was also aligned using the ClustaW tool to determine the conserved amino acid residues between the two proteins. It was observed that the crucial amino acids residues were conserved between the two proteins as seen in **Supplementary Figure S1B**. The amino acids involved in PAL (peptidoglycan associate lipoprotein) domain which aid in proper binding to the peptidoglycan is conserved in both, these being Asparagine (N68), Aspartic acid (D102), and Glycine (G105) [31, 32]. There is also the presence of an N-terminal, 13 amino acid long, linker sequence containing conserved critical residues. This linker sequence was predicted to be involved in binding to PAL domain, signaling carried out by YfiB, outer membrane sequestration of YfiR, and affects biofilm formation and adhesion of bacteria [28–32]. An important amino acid cluster from position 35 to 55 is present in the YfiB protein which is involved in the activation of YfiB and plays a role in appropriate YfiR sequestration by YfiB in the periplasm [28–32]. Identifying these important amino acids helped in selecting the targets for the following site-directed mutagenesis experiments.

To determine the 3D structure of *S. flexneri’*s YfiB protein, the I-TASSER tool was used, which predicts protein structure and function based on protein sequence, it identifies structural templates from the database by the multiple threading method known as LOMETS [61, 62]. Functional understanding of the target sequence is then attained by re-threading the 3D models using BioLiP, which is a protein functional database [61, 62]. To establish the structure of *Shigella* YfiB protein, I-TASSER utilized the solved crystal structure of YfiB protein from *Pseudomonas* [29, 30]. It was found that the YfiB protein is composed of a core OmpA-like domain (27-160 amino acids), has an N-terminal signal peptide (1-26 amino acids), and is comprised of α1-3 (three alpha-helices) and a β1-β4-β2-β3 network (anti-parallel β-sheet topology) (**Supplementary Figure S1C**).

Homologs of YfiB (E-value<10−4) in other gram-negative bacteria were identified and selected for determining the sequence conservation at each position and phylogenetic analysis. Conservation of amino acids at each position of the protein was analyzed using the WebLogo tool, all the YfiB homolog sequences were first aligned using ClustalW and then a sequence logo was created. **Supplementary Figure S2A** shows the graphical representation of all amino acids, the altitude of the pile at each position indicates sequence conservation and the height of the characters describes the comparative frequency of individual amino acids at that locus [63]. Using the ClustalW alignment of the homologs (**Supplementary Figure S2B**), an illustrative phylogenetic tree was created using the MEGA software [64], showing the evolutionary distance between the YfiB protein homologs from different gram-negative bacteria, calculated by the Maximum Composite Likelihood method [65, 66] (**Supplementary Figure S2C**).

### Mutational analysis of YfiB protein-Biofilm formation and Bacterial Adhesion Assays

To determine whether some of the conserved amino acid residues of YfiB are involved in the proper functioning of the protein, targeted mutagenesis was carried out. The targets were selected based on previous studies in *Pseudomonas* YfiBNR proteins and the in-silico analysis of YfiB protein in *S. flexneri* [29, 30]. It was found that the 13 amino acids long linker sequence was the most critical for YfiB functioning, as it helps in binding to the PAL domain, YfiB signaling, and sequestering the YfiR protein. In *Shigella*, the linker sequence is only 11 amino acids long, with the majority of amino acids being conserved. Three pairs of conserved amino acid residues from the linker sequence were selected as targets, these being Cys19Gln20 mutated to Ala19Glu20; Pro22Gln23 mutated to Ala22Glu23; Glu29Gln30 mutated to Ala29Glu30 (**Figure 7A**). Another conserved amino acid serine at position 36, which is part of an important amino acid cluster (35-55) involved in the activation and functioning of YfiB, was selected as a target and was mutated to alanine (**Figure 7B**). Mutants were cloned into pBAD_Myc_HisA plasmid and the protein expression of these mutants was also confirmed using western blot via an anti-HisA antibody to check if the amino acid mutations prevented the YfiB protein to be expressed properly (**Figure 7A & 7B; Supplementary Figure S3A & S3B**). To examine the effect of these mutations, biofilm formation assay and bacterial adhesion assay were chosen as the functional tests to see if the amino acid mutation caused alterations in YfiB activity and to the overall surface attachment and biofilm formation of the bacteria. These assays were performed, as explained in the methods section.

**Figure 7.**
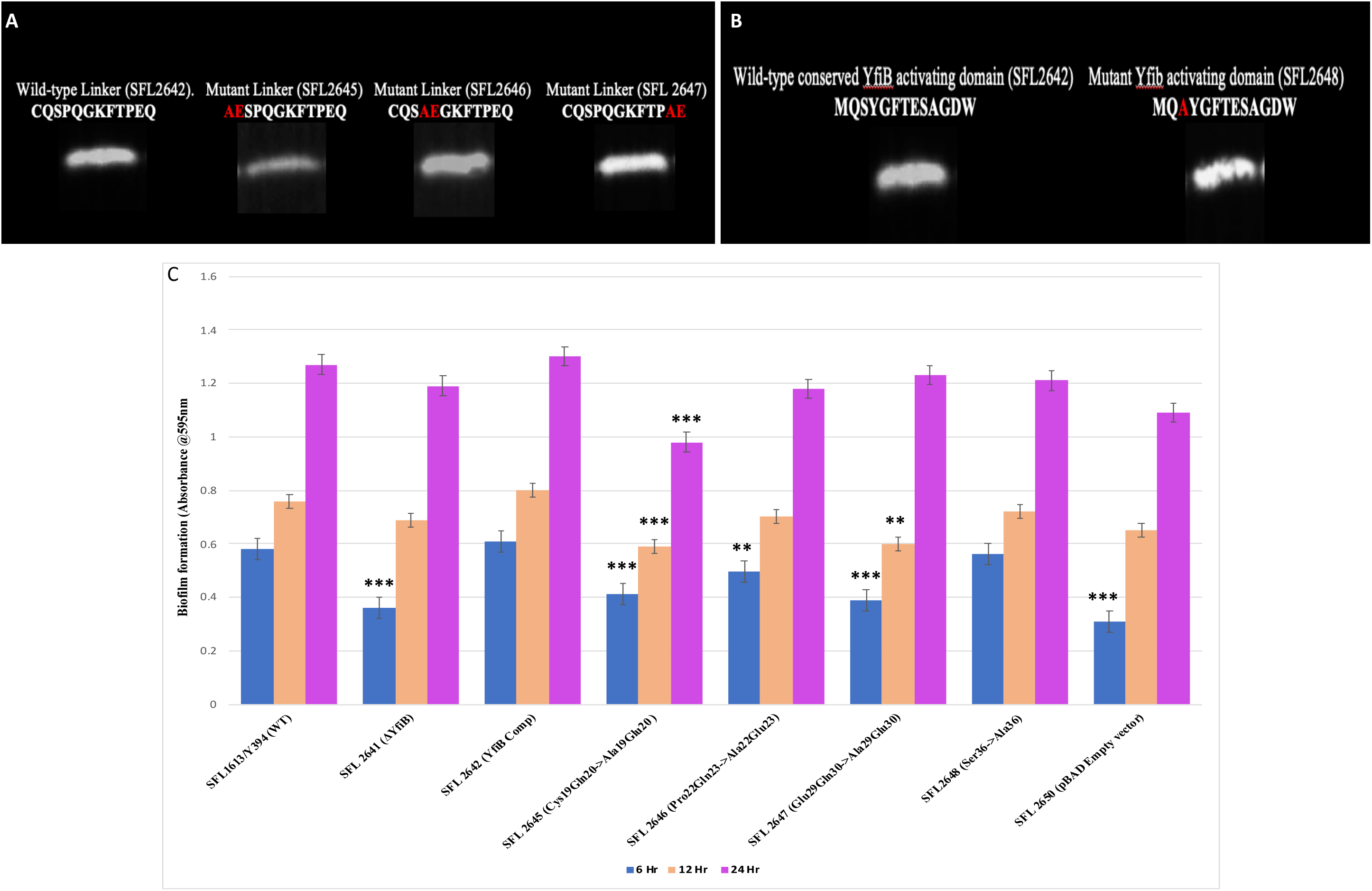
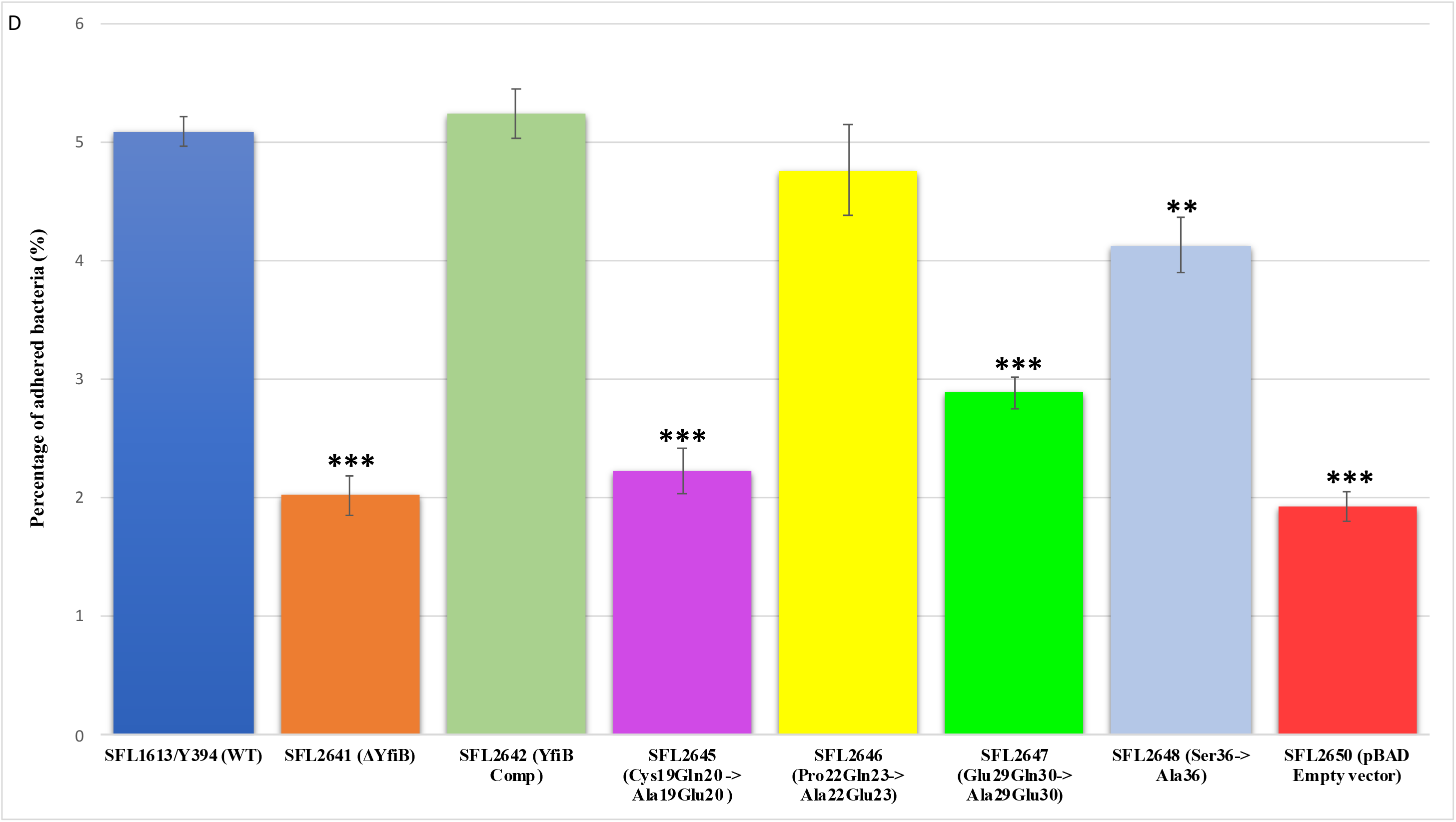
Analysing the effect of YfiB linker mutants and Serine36 mutant on YfiB activation and function. **(A)** The sequence of the wildtype N-terminal linker and the three site-directed linker mutants, the amino acid pairs mutated are highlighted in red. It also shows the immunoblot of the wild-type and mutant YfiB proteins in whole-cell lysate carried out with anti-HisA antibody. **(B)** The sequence of the wildtype YfiB activating domain with serine at position 36 and the mutant sequence with alanine at position 36, highlighted in red. It also shows the immunoblot of the wild-type and mutant YfiB proteins in whole-cell lysate carried out with anti-HisA antibody. **(C)** The consequence of various YfiB linker mutants and Ser36 mutant cloned in pBAD_Myc_HisA vector on biofilm formation, shown relative to the strain SFL2641/ΔYfiB containing the empty pBAD_Myc_HisA vector (SFL2650), used as control. The variance seen in the biofilm production between the strains was statistically evaluated using a student’s t-test. SFL2645 (Cys19Gln20->Ala19Glu20) mutant showed a decrease in biofilm at all three time points, whereas the SFL2646 (Pro22Gln23->Ala22Glu23) and SFL2647 (Glu29Gln30->Ala29Glu30) mutants only showed a decrease in observed biofilm at 6 and 12 hours. SFL2648 (Ser36 ->Ala36) showed only a slight decrease in biofilm formation when compared to the wildtype YfiB, but the difference wasn’t statistically significant. **(D)** The consequence of various YfiB linker mutants and Ser36 mutant cloned in pBAD_Myc_HisA vector on adherence to BHK cells, shown relative to the empty pBAD_Myc_HisA vector strain (SFL2650), used as control. To compute the variance seen in the percentage adhesion between the strains, a student’s t-test was performed. Mutants SFL2645 (Cys19Gln20->Ala19Glu20), SFL2647 (Glu29Gln30-> Ala29Glu30) and SFL2648 (Ser36->Ala36) showed a significant reduction in the percentage adhesion to BHK cells. Whereas mutant SFL2646 (Pro22Gln23->Ala22Glu23), showed a slight decrease when compared to the wildtype but the difference wasn’t substantial. Statistically significant (p. value < 0.05) variance is denoted by asterisks and the analysis was based on three independent repeats.

Biofilm formation assay was carried with the wild-type strain (SFL1613/Y394), *yfiB* deletion mutant (SFL2641/ΔYfiB) and the *yfiB* complemented strain (SFL2642/YfiBComp); along with the YfiB site-directed mutant strains-SFL2645 (Cys19Gln20->Ala19Glu20), SFL2646 (Pro22Gln23->Ala22Glu23), SFL2647 (Glu29Gln30->Ala29Glu30), and SFL2648 (Ser36->Ala36). The empty pBAD_Myc_HisA vector strain SFL2650 (SFL2641/ΔYfiB strain containing the empty pBAD_Myc_HisA plasmid) was used as the negative control. The assay was performed at 6, 12, and 24 hours and it was observed that there was a noteworthy difference between the biofilm formation of the wild-type YfiB protein and the functional mutants of YfiB (**Figure 7C**). SFL2645 (Cys19Gln20->Ala19Glu20) mutant showed a decrease in biofilm at all three-time points, whereas the SFL2646 (Pro22Gln23->Ala22Glu23) and SFL2647 (Glu29Gln30->Ala29Glu30) mutants only showed a decrease in observed biofilm at 6 and 12 hours. The mutant SFL2648 (Ser36->Ala36) showed only a slight decrease when compared to the wildtype YfiB strain, but the difference wasn’t statistically significant (**Figure 7C**).

Bacterial adhesion assay was carried out using the BHK cell line, as described in the methods. We infected the BHK monolayers with the strains SFL1613/Y394 (WT), SFL2641/ΔYfiB, and SFL2642/YfiBComp; along with the YfiB site-directed mutant strains-SFL2645 (Cys19Gln20->Ala19Glu20), SFL2646 (Pro22Gln23->Ala22Glu23), SFL2647 (Glu29Gln30->Ala29Glu30), and SFL2648 (Ser36->Ala36), and the empty pBAD_Myc_HisA vector strain (SFL2650). It was observed that the YfiB site-directed mutants showed a significantly reduced capacity to adhere to the epithelial cells of the host when compared with the wild-type YfiB protein. Mutants SFL2645 (Cys19Gln20->Ala19Glu20), SFL2647 (Glu29Gln30->Ala29Glu30) and SFL2648 (Ser36->Ala36) showed a significant reduction in the percentage adhesion to BHK cells. Whereas mutant SFL2646 (Pro22Gln23->Ala22Glu23), showed a slight decrease when compared to the wildtype but the difference wasn’t statistically significant (**Figure 7D**). It can be concluded from these functional assays that the conserved amino acid pairs in the linker sequence (Cys19Gln20, Pro22Gln23, and Glu29Gln30) and the serine at position 36, part of an important amino acid cluster of YfiB (35-55) are assumingly crucial for the activation and proper functioning of the YfiB protein.

## DISCUSSION AND CONCLUSION

In this work, we remarkably show that altering intracellular c-di-GMP levels can severely affect the pathogenesis of *Shigella flexneri*. This study suggests that the loss of outer-membrane YfiB protein, hinders the function of the inner membrane-bound YfiN (DGC activity), as the periplasmic YfiR is always bound to it, which causes a decreased concentration of c-di-GMP. Loss of YfiB and the apparent decrease in intracellular c-di-GMP levels, subsequently affect other downstream virulence factors of the bacteria, as seen by a slower biofilm production; decreased adhesion, and invasion of host cells; weakened ability to form plaques; and lower accumulation in guts of *C. elegans*, which leads to a surge in the life span of the worms. YfiB acts as a positive regulator of YfiN (a membrane-bound diguanylate cyclase), accountable for the production of c-di-GMP [70]. When YfiB is absent, it cannot sequester YfiR (acting as the negative regulator of YfiN) to the outer membrane and as a result, YfiN is always bound to YfiR. This causes YfiN to remain in an inactivated state, not being able to produce c-di-GMP and affecting various virulence functions in *Shigella* [28, 31, 32, 71]

Even though biofilm production in bacteria is dependent on a multitude of cellular factors and coordinated pathways, the slow biofilm production was seen in the *yfiB* deletion mutant can be explained by the functioning of the *yfiBNR* operon model, previously studied in *P. aeruginosa* [26, 27]. Disrupting the YfiBNR system by deleting the *yfiB* gene leads to a decreased c-di-GMP level, needed by *Shigella* for effective biofilm production in the presence of bile salts [19, 21, 37]. Higher intracellular levels of c-di-GMP are known to be responsible for an escalation of biofilm production and motility in bacteria [21–27]. Though in *E. coli*, it was observed that deleting genes encoding diguanylate cyclases such as *yfiN,* causing lower c-di-GMP levels, leads to higher motility and increased early biofilm production [70]. Motility affects biofilm formation as it has a functional role in preliminary surface attachment and attachment to the neighboring bacterial cells [21, 22, 70]. Bacterial motility also affects the architecture of biofilms, as it is known that strains with reduced motility produce flat biofilms while strains with prominent motility form vertical biofilm structures [14, 21]. This increased early biofilm formation observed in *E. coli* is not similarly observed in *Shigella*, presumably because they are non-motile bacteria and the decline in c-di-GMP directly decreases the biofilm formation. Earlier experiments in *E. coli* have proven that non-flagellated or bacterial cells with paralyzed flagella show decreased biofilm production in a lower concentration of c-di-GMP [70, 72], which confirms the observed slow biofilm production in *Shigella-yfiB* deletion mutant.

The consequence of *yfiB* deletion on *Shigella’*s ability to adhere, invade and spread in the host cells can also be explained by the decreased intracellular levels of c-di-GMP in the deletion mutant. c-di-GMP signaling is known to be interrelated to various bacterial virulence phenotypes like host cell invasion, secretion of virulence factors, cytotoxicity, intracellular infection, host-cell adherence, cell motility, alteration of immune responses, and resistance to oxidative stress [21–27]. Invasion and injecting bacterial effectors into the host cells which is mediated by the type 3 secretion system in *Shigella,* is also affected by the cellular concentration of c-di-GMP [22, 73]. High concentrations of c-di-GMP are known to have a progressive effect on the adhesion and invasion of the bacteria which eventually aids the pathogen in colonizing the host epithelial cells [21–27]. This relationship between biotic/abiotic surface attachment and c-di-GMP concentration has been previously inspected in the opportunistic human pathogen *P. aeruginosa* [31, 32, 74] and recently in *S. flexneri* [20]. It is known from these studies, that on surface contact, there is an upsurge in levels of c-di-GMP, which induces flagella/pilli biosynthesis, surface adherence and boosts virulence [22]. This explains how by deleting the *yfiB* gene and obstructing the proper functioning of the *yfiBNR* operon, we can severely affect the pathogenicity of *Shigella* by impeding its adhesion, invasion, and intercellular spreading capabilities.

The *C. elegans* accumulation assay to test changes in *Shigella* virulence after deleting *yfiB* gene also validated the same that YfiB acts as a positive regulator of YfiN producing c-di-GMP. Low accumulation of *yfiB* deletion mutant in the *C. elegans* intestinal lumen, indicates that it becomes less virulent, eventually increasing the lifespan and survival of worms in comparison to the wild-type strain. This shows how YfiB and the YfiBNR system have a significant impact on the intestinal accumulation of *S. flexneri* in the guts of *C. elegans* worms. It is known that several bacterial species and non-virulent bacteria are unable to persistently colonize the nematode gut, are rapidly digested and expelled from the intestine [46, 47, 75]. c-di-GMP catabolizing enzymes have been previously shown, supporting intestinal gut colonization by the bacteria and improving its intracellular survival [76, 77]. As discussed earlier, c-di-GMP plays a big role in bacterial adherence and invasion processes, which are necessary steps to invade the *C. elegans* gut lumen [21–27]. Hence the low bacterial accumulation seen of the *yfiB* deletion mutant in *C. elegans* gut sits consistent with the other observations of a slow biofilm production; reduced adherence and invasion; and decreased plaque numbers; when there is decreased c-di-GMP concentration inside the cells owing to the disruption of *yfiBNR* operon caused by the deletion of the *yfiB* gene.

Mutational studies of the YfiB protein in this study showed how the conserved amino acid pairs of the linker sequence and the conserved serine at position 36 are essential for its proper functioning. Protein expression of the mutants was also checked via western blotting of whole-cell lysates to confirm if the mutants caused any expression defects. The slight differences that were seen in the proteins bands of the site-directed mutants even after equal protein loading may be due to the affected expression of YfiB caused by the amino acid mutations or due to an experimental artifact. The linker sequence acts as a lipid anchor and helps in peptidoglycan binding, which is critical for the full activity of the YfiB [29, 30]. Anchoring of YfiB in the cell wall and outer membrane aids in proper signaling and accurate YfiR sequestration, which eventually activates the YfiN, leading to c-di-GMP production [29–32]. Amino acid mutation of the linker sequence can result in decreased c-di-GMP levels due to improper functioning of the YfiBNR system [29, 30]. This can have a significant effect on all downstream virulence-related progressions in bacteria such as the formation of biofilm, attachment, and invasion of the host cells to name a few which are regulated by c-di-GMP [29–32]. The serine at position 36 was also found to be conserved in most YfiB homologs and had the highest frequency of being present at this position when compared to other similar amino acids (**Supplementary Figure S2A**). This was mutated to alanine to test its function (SFL 2648/Ser36->Ala36) and it was observed that it doesn’t exhibit any difference in the biofilm production but demonstrates a significant decrease in the percentage adhesion to BHK cells when compared to the wildtype YfiB. Serine at position 36 is part of an important amino acid cluster of YfiB (35-55) which is known to be involved in the activation of YfiB protein, proper YfiR sequestration, and effects c-di-GMP production [29, 30]. This functional analysis proves that YfiN is under positive and negative control of YfiB and YfiR respectively and together this YfiBNR system regulates the intracellular quantities of c-di-GMP, which aids in various virulence mechanisms of the bacteria.

Each year, *Shigella* is known to cause millions of infections globally and has become a major public health concern. There is a gradual decrease of treatment alternatives due to the rise in antimicrobial resistance and a lack of *Shigella* vaccine which could prevent infections [1–9]. Hence it has become extremely important to understand every aspect of the *Shigella* infection model, to develop newer drug targets and possible vaccine candidates. This present study establishes that the c-di-GMP regulation by the YfiBNR system is involved in *S. flexneri’s in-vivo* persistence, biofilm formation, and its infection cycle. c-di-GMP is a well-known player in bacterial virulence, promoting prolonged survival of the pathogen and as homologs of YfiBNR exist in other gram-negatives, this regulatory system can be used as an effective drug target in treating bacterial diseases [78–80]. Antibacterial drugs that can affect the functioning of YfiB or YfiN directly and/or other diguanylate cyclases present in the bacteria can be used effectively to control the spread and decrease the span of bacterial infections. This also reduces our dependency on antimicrobials to treat infections and the risk of generating any multidrug-resistant strains. This study provides a start point for mechanistic and functional research of the YfiBNR signaling system and c-di-GMP mediated virulence in *Shigella*. Additionally inspecting the prospect of targeting this signaling network, for discovering new therapeutics for bacterial infections like shigellosis or the ones caused by other bacterial pathogens.

## Author Information

### Affiliations

**Division of Biomedical Science and Biochemistry, Research School of Biology, The Australian National University, ACT, Canberra, Australia.** Tanuka Sen & Naresh K.Verma.

### Contributions

T.S. - designed and conducted the experiments; analysed and interpretation of data; prepared figures and wrote the manuscript. N.K.V.-conceived and directed the study and critically revised the manuscript. Both the authors read and approved the final manuscript.

## Corresponding author

Correspondence to Naresh K. Verma.

## Availability of supporting data

N/A. Additional data is provided in the supplementary file.

## Supplementary material

Supplemental data for this article- Supplementary zip file.

