## Supplemental Tables and Figure for "YfiB: An Outer Membrane Protein Involved in the Virulence of *Shigella flexneri*": Supplementary Figures.pdf

**This file includes Supplementary Figures S1, S2 and S3.**

**Figure S1- Results for conserved domain analysis; predicted protein structure by I-TASSER and ClustalW alignment of YfiB from *Pseudomonas* and *Shigella*. (A)** Conserved domains found in the YfiB protein, analysed using NCBI-protein BLAST and conserved domain database (CDD). **(B)** ClustalW protein sequence alignment of YfiB protein from *Pseudomonas and Shigella*, showing highly/weakly similar and dissimilar amino acids; it also highlights the conserved essential domains. *Blue-conserved linker sequence; Purple- 35-55 amino acid cluster important for activating YfiB; Yellow - PAL Domain associated amino acids. **(C)** The 3D protein structure predicted for YfiB, computed by the I-TASSER tool, using *Pseudomonas* YfiB solved crystal structure as a template; the 3D structure consists of Core OmpA-like domain, four-stranded anti-parallel β-sheet with topology β1-β4-β2-β3 and helices α1-3

**A**

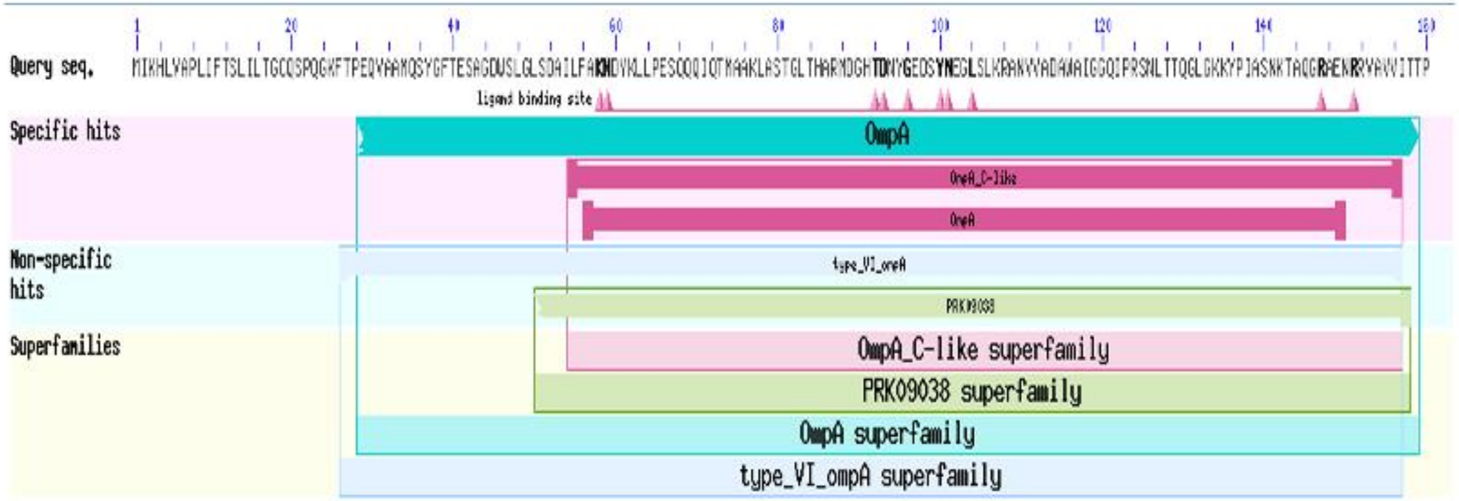


**B**

```
YfiB_Pseudomonas    MLPQRLHPSRLLALALFSLVLGLAGCQTKPPQTGLSAEQIAVLQEQGFELRDEGWEFGMS 60
YfiB_Shigella       MIKH-------LVAPLIFTSLILTGCQ--SPQGKFTPEQVAAMQSYGFTESAGDWSLGLS 51
                    *: :       *.  *:    * *:***    **   :: **:*.:*. **     .*.:*:*

YfiB_Pseudomonas    SKVLFGNNLDRLNPDSRNTLTKIARALLAVDIDKVRLEGHTDNYGDEGYNQKLSERRAES 120
YfiB_Shigella       DAILFAKNDYKLLPESQQQIQTMAAKLASTGLTHARMDGHTDNYGEDSYNEGLSLKRANV 111
                    . :**.:*  :* *:*:: : .:*  * :..: :.*::*******::.**: ** :**:

YfiB_Pseudomonas    VAAVFREA-GMPAANIEVRGLGMSKPVADNKTRAGRSENRRVAIIVPAE   168
YfiB_Shigella       VADAWAIGGQIPRSNLTTQGLGKKYPIASNKTAQGRAENRRVAVVITTP   160
                    ** .:  .  :* :*: .:*** . *:*.***  **:******::: :
```

**C**

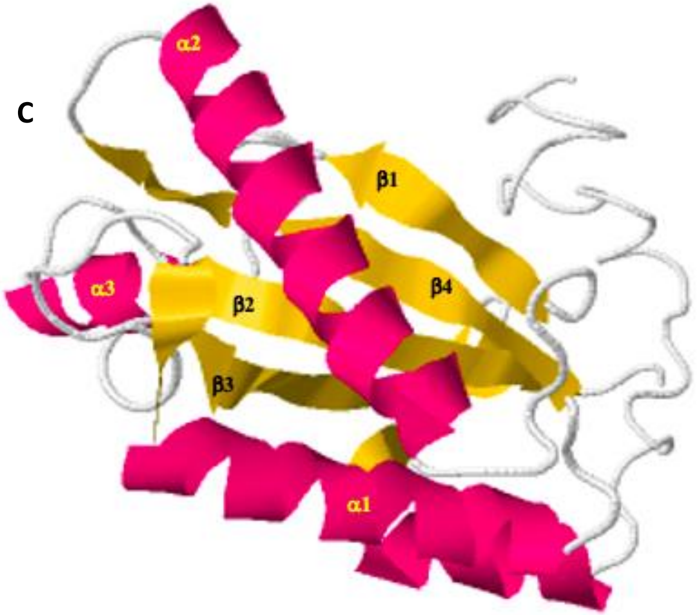

**Figure S2- In-silico investigation of the YfiB protein. (A)** Weblogo representations of the YfiB protein amino acid conservation. The height of amino acid code at each position reflects the comparative incidence of the amino acid at that position and the total height of the pile denotes the degree of conservation at each individual locus (measured in bits). The consensus sequence was derived from multiple sequence alignment of YfiB protein sequences from various gram-negative bacteria. **(B)** ClustalW multiple sequence alignment of YfiB protein homologs found in various gram-negative bacteria, showing conserved and non-conserved amino acid residues. **(C)** Phylogenetic tree illustrating the evolutionary distance between the YfiB protein homologs from various gram-negative bacteria. ClustalW alignment of the homologs was used to generate this phylogenetic tree and was created using the MEGA software, calculated by the Maximum Composite Likelihood method.

**A**

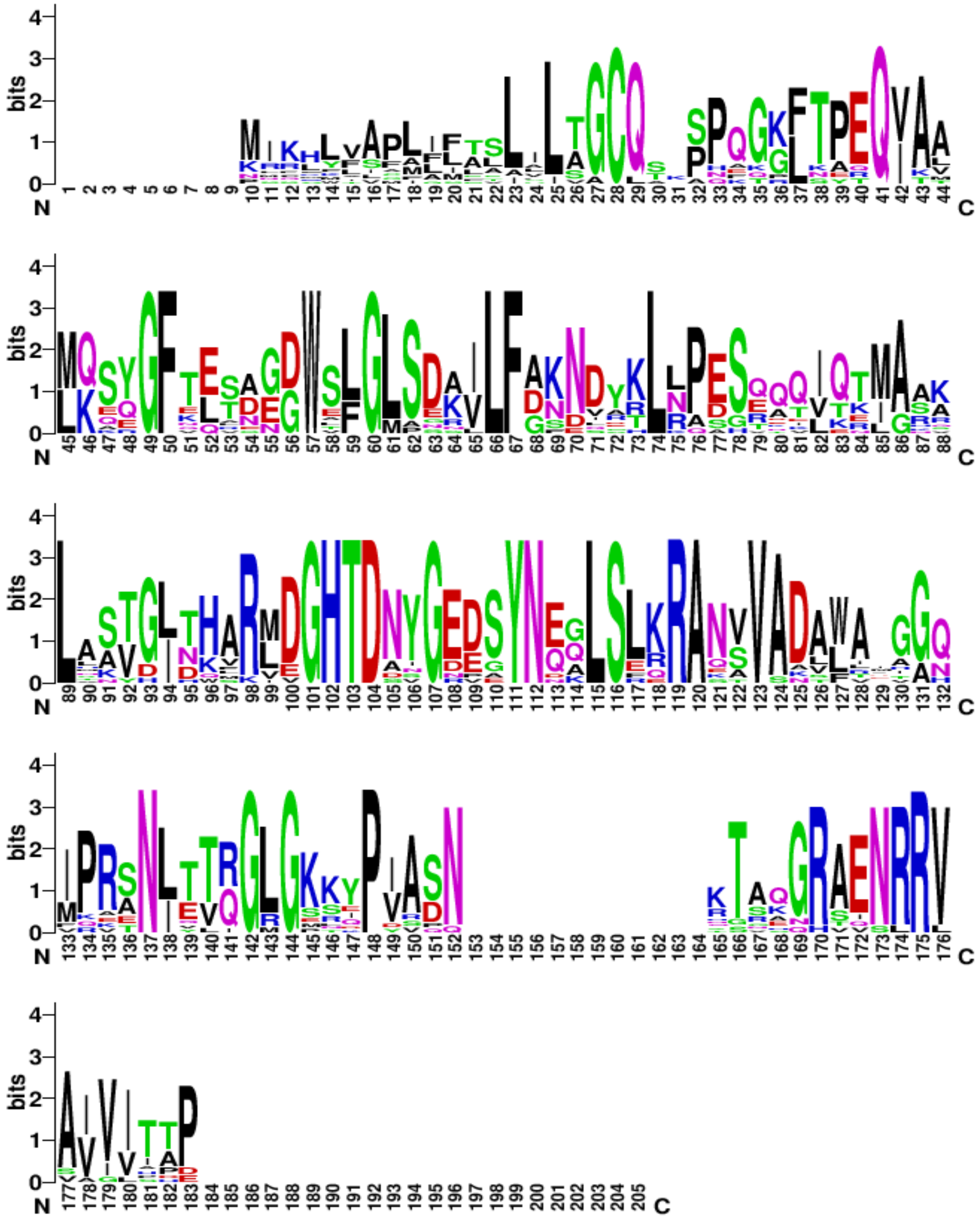

**Figure S2- In-silico investigation of the YfiB protein. (A)** Weblogo representations of the YfiB protein amino acid conservation. The height of amino acid code at each position reflects the comparative incidence of the amino acid at that position and the total height of the pile denotes the degree of conservation at each individual locus (measured in bits). The consensus sequence was derived from multiple sequence alignment of YfiB protein sequences from various gram-negative bacteria. **(B)** ClustalW multiple sequence alignment of YfiB protein homologs found in various gram-negative bacteria, showing conserved and non-conserved amino acid residues. **(C)** Phylogenetic tree illustrating the evolutionary distance between the YfiB protein homologs from various gram-negative bacteria. ClustalW alignment of the homologs was used to generate this phylogenetic tree and was created using the MEGA software, calculated by the Maximum Composite Likelihood method.

**B**

```
YfiB_Citrobacter                    --------------------------------------------MKSYGF
YfiB_Enterobacter                   ---------MLKRYFAPLLLASLVISGCQT-SPEGKFTPEQIAAMKSYGF
YfiB_E.coli                         ---------MIKHLVAPLIFTSLILTGCQ--SPQGKFTPEQVAAMQSYGF
YfiB_S.dysenteriae                  ---------MIKHLVAPLIFTSLILTGCQ--SPQGKFTPEQVAAMQSYGF
YfiB_S.flexneri_X                   ---------MIKHLVAPLIFTSLILTGCQ--SPQGKFTPEQVAAMQSYGF
YfiB_S.sonnei                       ---------MIKHLVAPLIFTSLILTGCQ--SPQGKFTPEQVAAMQSYGF
YfiB_SFL1613/Y394                   ---------MIKHLVAPLIFTSLILTGCQ--SPQGKFTPEQVAAMQSYGF
YfiB_S.boydii                       --------------------------------------------MQSYGF
YfiB_Klebsiella_pneumoniae          ---------MIRKYFVPALMAAALLTGCQ--APQGKFTPEQVAAMKSYGF
YfiB_Xenorhabdus                    ------MSKKRSFMFICAFIGTLFLSACQ---NKGGLTAEQITTLKQQGF
YfiB_Pseudomonas_chlororaphis       ----------MQLFTAGLLLALLSLTGCQSVPP-KGLTPQQVAVLKQEGF
YfiB_Pseudomonas_aeruginosa         --MLPQRLHPSRLLALALFSLVLGLAGCQTKPPQTGLSAEQIAVLQEQGF
YfiB_Yersinia_enterocolitica        MLGLNNNRQKNPLWISFFALCLLVLVGCQAKPH-QGLTPEQIAALQEQGF
YfiB_Serratia                       --MIQQTLKNRFSLLAMMFIALLALAGCQSKPQ--GLTPEQIALLQSQGF
YfiB_Aeromonas                      -----------------MGMFALLLAGCQ-SPPAGRLNETQIALLKARGF
YfiB_Acinetobacter_baumannii        ------------MKLSFIALLCIALAGCLS---FGPLKYRQVKMLKKEGF
                                                                                ::   **

YfiB_Citrobacter                    TEASGDWSLGLSDNILFDKNDYKLRPESEKQIKEMASKLAATGLNHARLD
YfiB_Enterobacter                   NELNGDWSLGLSDKILFDKNDARLRPESETQIQTMASRLAATGLNHARMD
YfiB_E.coli                         TESAGDWSLGLSDAILFAKNDYKLLPESQQQIQTMAAKLASTGLTHARMD
YfiB_S.dysenteriae                  TESAGDWSLGLSDAILFAKNDYKLLPESQQQIQTMAAKLASTGLTHARMD
YfiB_S.flexneri_X                   TESAGDWSLGLSDAILFAKNDYKLLPESQQQIQTMAAKLASTGLTHARMD
YfiB_S.sonnei                       TESAGDWSLGLSDAILFAKNDYKLLPESQQQIQTMAAKLASTGLTHARMD
YfiB_SFL1613/Y394                   TESAGDWSLGLSDAILFAKNDYKLLPESQQQIQTMAAKLASTGLTHARMD
YfiB_S.boydii                       TESAGDWSLGLSDAILFAKNDYKLLPESQQQIQTMAAKLTSTGLTHARMD
YfiB_Klebsiella_pneumoniae          TESNGDWSLGLSDSILFDKNDYRLRPDSRQQITTMASRLAATGITHSRLE
YfiB_Xenorhabdus                    QQTDEGWLFGMSEKVLFGNNQSHLRPEGEAKLKELASVLSKVGIHHARLD
YfiB_Pseudomonas_chlororaphis       ELTDEGWAFGLSGKVLFGSDIETLNAASTEIVERIGKALVSVHIDKVRVD
YfiB_Pseudomonas_aeruginosa         ELRDEGWEFGMSSKVLFGNNLDRLNPDSRNTLTKIARALLAVDIDKVRLE
YfiB_Yersinia_enterocolitica        KLTDNGWVFGLANKVLFDSDVRKLNASGVQTVQNIGRALHNVGINHMRVD
YfiB_Serratia                       KLTDNGWEFGLSDKVLFGNNVGKLNPESTDTVQKMGRALMSVGITQFRLD
YfiB_Aeromonas                      TQVEEGWSLGLSDRVLFANESRTLNPDSRAVVQKIAHSLLSVGIDWARLD
YfiB_Acinetobacter_baumannii        VLTNEGWTLGLPERLLFDFNDATLKQSHEAELTRLANQLNKYDLNKLKIV
                                         .* :*:.  :**  :   *       :  :.  *    :   ::

YfiB_Citrobacter                    GHTDNYGEDSYNEALSLKRANVVADAWAQGANIPRTNLTTQGLGKKYPIA
YfiB_Enterobacter                   GHTDNYGEESYNEALSLKRANVVADAWAKGANIPRSNLTTRGLGKKYPVS
YfiB_E.coli                         GHTDNYGEDSYNEGLSLKRANVVADAWAMGGQIPRSNLTTQGLGKKYPIA
YfiB_S.dysenteriae                  GHTDNYGEDSYNEGLSLKRANVVADAWAMGGQIPRSNLTTQGLGKNIP--
YfiB_S.flexneri_X                   GHTDNYGEDSYNEGLSLKRANVVADAWAIGGQIPRSNLTTQGLGKKYPIA
YfiB_S.sonnei                       GHTDNYGEDSYNEGLSLKRANVVADAWAIGGQIPRSNLTTQGLGKKYPIA
YfiB_SFL1613/Y394                   GHTDNYGEDSYNEGLSLKRANVVADAWAIGGQIPRSNLTTQGLGKKYPIA
YfiB_S.boydii                       GHTDNYGEDSYNEGLSLKRANVVADAWAMGGQIPRSNLTTQGLGKKYPIA
YfiB_Klebsiella_pneumoniae          GHTDNYGEDSYNEALSLKRANSVADAWAEGAHVPRSNLVTRGLGKKIPDR
YfiB_Xenorhabdus                    GHTDNYGEVSYNNQLSLKRANTVADALTDGG-MQRANLTTRGLGPSQPIA
YfiB_Pseudomonas_chlororaphis       GHTDASGREAYNQQLSLRRAKSVSKVLVATG-MREENIQLRGLGSSEPVA
YfiB_Pseudomonas_aeruginosa         GHTDNYGDEGYNQKLSERRAESVAAVFREAG-MPAANIEVRGLGMSKPVA
YfiB_Yersinia_enterocolitica        GHTDAIGEDGYNQQLSFQRASAVADTLAAIG-IPRTNIEVRGRGKLEPVA
YfiB_Serratia                       GHTDNYGEDSYNDQLSLRRANAVADLLASVG-IPRANIETRGMGKRDPVA
YfiB_Aeromonas                      GHTDSNGDESYNQQLSRQRAQSVADSLIDAG-MPVANLEVRGLGERYPVA
YfiB_Acinetobacter_baumannii        GHTDDVGNPEYNQKLSEERAQSVANLFLTHG-FKKENIYVIGRGSTQPYV
                                    ****  *   **: ** .**. *:      . .   *:   * *   *

YfiB_Citrobacter                    SN------------QTSKGRAENRRVAVVITTP-----------------
YfiB_Enterobacter                   SN------------HTAQGRAENRRVAVVISTP-----------------
YfiB_E.coli                         SN------------KTAQGRAENRRVAVVITTP-----------------
YfiB_S.dysenteriae                  --------------------------------------------------
YfiB_S.flexneri_X                   SN------------KTAQGRAENRRVAVVIATP-----------------
YfiB_S.sonnei                       SN------------KTAQGRAENRRVAVVITTP-----------------
YfiB_SFL1613/Y394                   SN------------KTAQGRAENRRVAVVITTP-----------------
YfiB_S.boydii                       SN------------KTAQGRAENRRVAVVITTP-----------------
YfiB_Klebsiella_pneumoniae          QQRHGGRARRKPPGNGGHQHAVSLRLVAGLHHPHCRCRAPAQVVNGNEQQ
YfiB_Xenorhabdus                    DN------------RSSKGRAENRRVAIVITAP-----------------
YfiB_Pseudomonas_chlororaphis       SN------------STVAGRTENRRVSIVVIAD-----------------
YfiB_Pseudomonas_aeruginosa         DN------------KTRAGRSENRRVAIIVPAE-----------------
YfiB_Yersinia_enterocolitica        DN------------RTAKGRAENRRVAIIVTAP-----------------
YfiB_Serratia                       DN------------RTSSGRVENRRVAIVVTP------------------
YfiB_Aeromonas                      SN------------KTREGRSQNRRVAIVITD------------------
YfiB_Acinetobacter_baumannii        PN------------TTNENRAINRRVAIVIIP------------------

YfiB_Citrobacter                    -----
YfiB_Enterobacter                   -----
YfiB_E.coli                         -----
YfiB_S.dysenteriae                  -----
YfiB_S.flexneri_X                   -----
YfiB_S.sonnei                       -----
YfiB_SFL1613/Y394                   -----
YfiB_S.boydii                       -----
YfiB_Klebsiella_pneumoniae          QAGSH
YfiB_Xenorhabdus                    -----
YfiB_Pseudomonas_chlororaphis       -----
YfiB_Pseudomonas_aeruginosa         -----
YfiB_Yersinia_enterocolitica        -----
YfiB_Serratia                       -----
YfiB_Aeromonas                      -----
YfiB_Acinetobacter_baumannii        -----
```

**Figure S2- In-silico investigation of the YfiB protein. (A)** Weblogo representations of the YfiB protein amino acid conservation. The height of amino acid code at each position reflects the comparative incidence of the amino acid at that position and the total height of the pile denotes the degree of conservation at each individual locus (measured in bits). The consensus sequence was derived from multiple sequence alignment of YfiB protein sequences from various gram-negative bacteria. **(B)** ClustalW multiple sequence alignment of YfiB protein homologs found in various gram-negative bacteria, showing conserved and non-conserved amino acid residues. **(C)** Phylogenetic tree illustrating the evolutionary distance between the YfiB protein homologs from various gram-negative bacteria. ClustalW alignment of the homologs was used to generate this phylogenetic tree and was created using the MEGA software, calculated by the Maximum Composite Likelihood method.

**C**

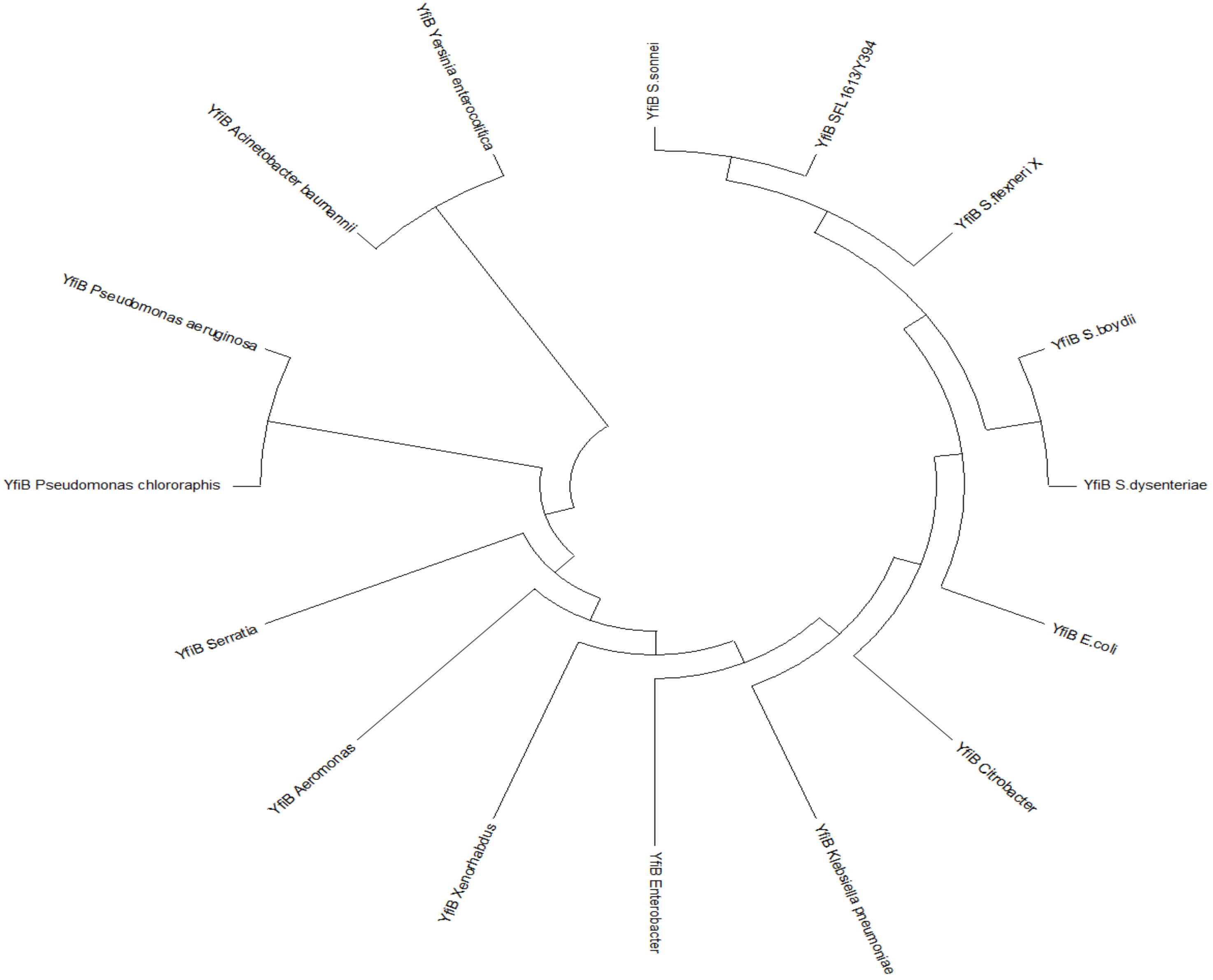

**Figure S3- Confirming protein expression of the YfiB wildtype protein and the YfiB site directed mutants. (A)** SDS gel stained with coomassie blue dye for validating equal total protein loading for each sample. **(B)** Western transfer and blotting of the loaded SDS gel using the anti-HisA antibody to check for correct protein expression. YfiB is known to be a 17.2 kDa protein, wildtype YfiB protein sample and site directed mutants of YfiB, all show a band at the same location. The intensity of the bands maybe due to an experimental artifact or an effect of amino acid mutation on protein expression. *Lane 1- Pre-stained protein ladder (10-250 kDa); Lane 2- SFL2650 (SFL2641/ΔYfiB strain containing the empty pBAD_Myc_HisA vector); Lane 3- Wildtype YfiB (SFL2642/YfiBComp); Lane 4- SFL2645 (Cys19Gln20->Ala19Glu20); Lane 5- SFL2646 (Pro22Gln23->Ala22Glu23); Lane 6- SFL2647 (Glu29Gln30->Ala29Glu30); & Lane 7- SFL2648 (Ser36->Ala36).

A

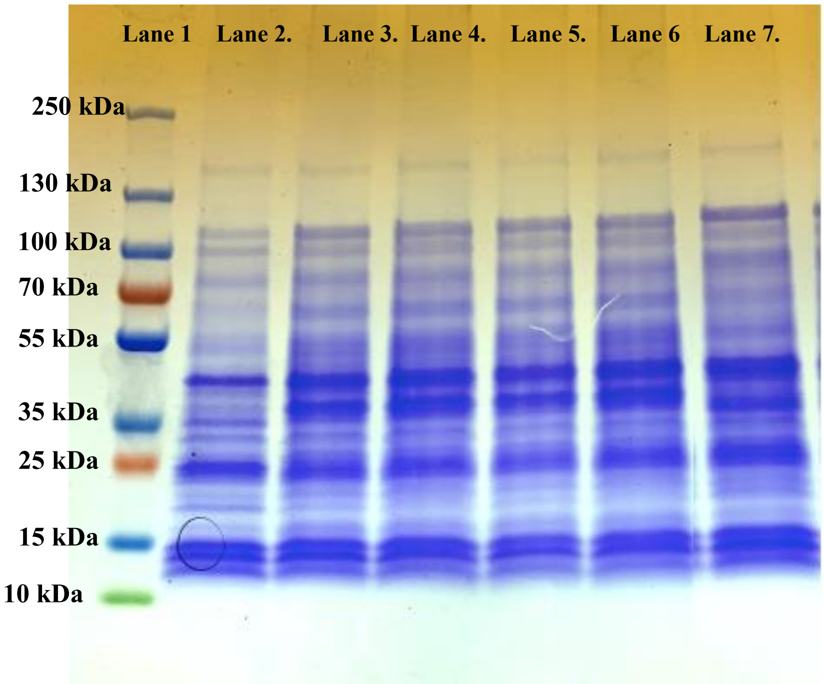


B

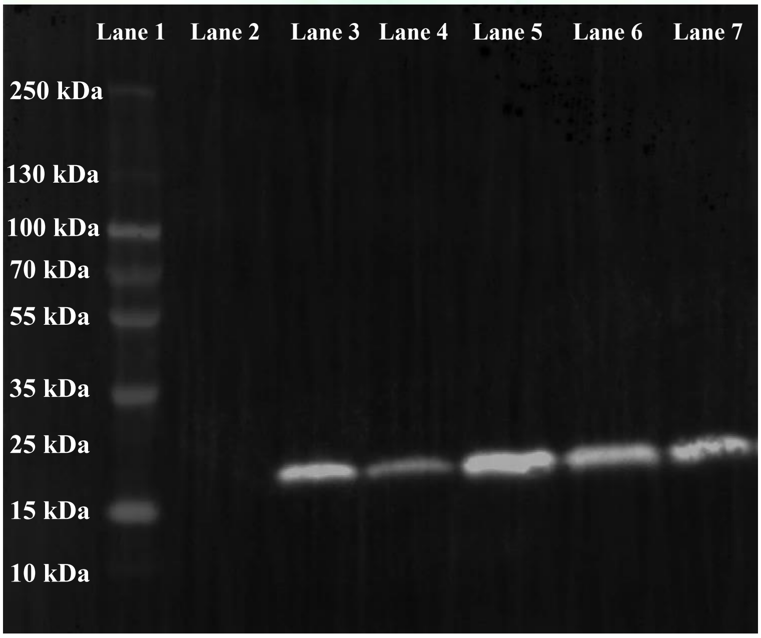
