## Supplemental Tables and Figure for "YfiB: An Outer Membrane Protein Involved in the Virulence of *Shigella flexneri*": Supplementary Tables_YfiB.pdf

This file includes Tables S1, S2, and S3.

**Table S1- Bacterial Strains and plasmids used in this study**

| Name | Description | Antibiotic Resistance |
|---|---|---|
| ***Shigella flexneri*** | | |
| SFL1613/Y394 | Wild-type *S.flexneri 1c strain*[26] | - |
| SFL2641/ΔYfiB | Y394 with *yfiB* deleted | CM |
| SFL2642/YfiBComp | SFL2641 with *yfiB* gene complemented, cloned into pBAD_Myc_HisA vector | Amp, Ery, CM |
| SFL2645 | SFL 2641 with site directed mutant- Cys19Gln20-> Ala19Glu20 , cloned into pBAD_Myc_HisA vector. | Amp, Ery, CM |
| SFL2646 | SFL 2641 with site directed mutant- Pro22Gln23-> Ala22Glu23, cloned into pBAD_Myc_HisA vector. | Amp, Ery, CM |
| SFL2647 | SFL 2641 with site directed mutant- Glu29Gln30 -> Ala29Glu30, cloned into pBAD_Myc_HisA vector. | Amp, Ery, CM |
| SFL2648 | SFL 2641 with site directed mutant- Ser36-> Ala36, cloned into pBAD_Myc_HisA vector. | Amp, Ery, CM |
| SFL2650 | SFL 2641 with empty pBAD_Myc_HIsA vector | Amp, Ery, CM |
| ***E. coli*** | | |
| B2298 | OP50 *E. coli* strain | - |
| **Plasmids** | | |
| pKD46 | Helper plasmid expressing the lambda red genes (*gam, beta*, and *exo*) | Amp |
| pkD3 | Used for template generation for homologous recombination. | Cm |
| pBAD_Myc_HisA | Vector used for cloning *yfiB* gene, used as control empty vector. | Amp, Ery |
| pBAD_Myc_HisA_YfiB | pBAD_Myc_HisA with *yfiB* gene cloned at the *Nco*I and *Hind*III. | Amp, Ery |
| pBAD_Myc_HisA_M1 | pBAD_Myc_HisA with mutant YfiB (Cys19Gln20 -> Ala19Gln20) | Amp, Ery |
| pBAD_Myc_HisA_M2 | pBAD_Myc_HisA with mutant YfiB (Pro22Gln23 -> Ala22Glu23) | Amp, Ery |
| pBAD_Myc_HisA_M3 | pBAD_Myc_HisA with mutant YfiB (Glu29Gln30 -> Ala29Glu30) | Amp, Ery |
| pBAD_Myc_HisA_M4 | pBAD_Myc_HisA with mutant YfiB ( Ser36-> Ala36) | Amp, Ery |

**Table S2 - Primers used in this study**

| Primer Name | Role | Sequence |
|---|---|---|
| *yfiB*_For | Primer for amplifying the CM gene of pKD3, containing the 80bp overhangs, homologous to the upstream of the *yfiB* gene | AAGAGCTTGCCGATCACAATATGTATCAGGCCAAACAC CAGCGTG<br>CCGAAAAGCTGGTGAGATAACAAGGATATATCGATTA CACGTCTT<br>GAGCGATTGT |
| *yfiB*_Rev | Primer for amplifying the CM gene of pKD3, containing the 80bp overhangs, homologous to the downstream of the *yfiB* gene | GGCTGCTCGTATCAAAGAGCGTCTTAACTAAGATTTCG CTTAAGCGA<br>CATCCTGTTAAGAAGGGCTGGCCAATTGGCTGGGGTCC ATATGAA<br>TATCCTCC |
| *yfiB*_Test_For | Confirming deletion of *yfiB* gene | AGCCGATTCGTCACAATGGT |
| *yfiB*_Test_Rev | Confirming deletion of *yfiB* gene | CTCCGGTAGTTGACAGCATT |
| CM_Test | Confirming deletion of *yfiB* gene and presence of chlorampheni col gene | CAACGGTGGTATATCCAGTG |
| Apy_For | Checking presence of *Shigella*'s Virulence plasmid | CATAATCAAGAGACAAAACGATA |
| Apy_Rev | Checking presence of *Shigella*'s Virulence plasmid | CCAGCCTTTCCAGTAATCCC |

| | | |
|---|---|---|
| VirG_For | Checking presence of *Shigella*'s Virulence plasmid | CGGGTACTCAAGAACTTCAAT |
| VirG_Rev | Checking presence of *Shigella*'s Virulence plasmid | TTCCGCCAAAATGAGAGTTCC |
| pBAD_Univ_For | Primer to check for correct cloning in the pBAD_Myc_HisA vector | ATGCCATAGCATTTTTATCC |
| pBAD_Uni_Rev | Primer to check for correct cloning in the pBAD_Myc_HisA vector | GATTTAATCTGTATCAGG |

**Table S3- UniProt accession number of the YfiB protein sequences used in this study for in-silico analysis.**

| UniProt ID[b] | Organism |
|---|---|
| C3SYR2 | *Escherichia coli* |
| A0A377W1F8 | *Klebsiella pneumoniae* |
| A0A2X2HQ42 | *Shigella boydii* |
| A0A2X2ISV4 | *Shigella dysenteriae* |
| D2AHI5 | *Shigella flexneri* serotype X (strain 2002017) |
| Q3YYN3 | *Shigella sonnei* (strain Ss046) |
| A0A2I8JE62 | *Shigella flexneri 1c* strain (Y394) |
| Q9I4L6 | *Pseudomonas aeruginosa (strain ATCC 15692)* |
| A0A3G7JRF7 | *Pseudomonas chlororaphis subsp. aurantiaca* |
| A0A7G2IWZ5 | *Citrobacter freundii* |
| A0A7U4NUK4 | *Acinetobacter baumannii* |
| A0A0S2SF47 | *Aeromonas schubertii* |
| G8LNV7 | *Enterobacter ludwigii* |
| N1NGC1 | *Xenorhabdus nematophila F1* |
| A0A1W5DTJ7 | *Serratia proteamaculans* |
| A0A485DSF5 | *Yersinia enterocolitica* |
